# Escape of TLR5 Recognition by *Leptospira spp*: A Rationale for Atypical Endoflagella

**DOI:** 10.1101/2020.05.30.121202

**Authors:** Marion Holzapfel, Delphine Bonhomme, Julie Cagliero, Frédérique Vernel-Pauillac, Martine Fanton d’Andon, Sophia Bortolussi, Laurence Fiette, Cyrille Goarant, Elsio A. Wunder, Mathieu Picardeau, Albert I. Ko, Dirk Werling, Mariko Matsui, Ivo G. Boneca, Catherine Werts

## Abstract

*Leptospira interrogans* are invasive bacteria responsible for leptospirosis, a worldwide zoonosis. They possess two periplasmic endoflagella that allow their motility. *L. interrogans* are stealth pathogens that escape the innate immune responses of the NOD-like receptors NOD1/2, and the human Toll-like receptor (TLR)4, sensing peptidoglycan and lipopolysaccharide (LPS), respectively. TLR5 is another receptor of bacterial cell wall components, recognizing flagellin subunits.

To study the contribution of TLR5 in the host defense against leptospires, we infected WT and TLR5 deficient mice with pathogenic *L. interrogans* and tracked the infection by *in vivo* live imaging of bioluminescent bacteria or by q-PCR. We did not identify any protective or inflammatory role of murine TLR5 to control pathogenic *Leptospira*. Likewise, subsequent *in vitro* experiments showed that infections with different live strains of *L. interrogans* and *L. biflexa* did not trigger TLR5. However, unexpectedly, heat-killed bacteria stimulated human and bovine TLR5, although barely mouse TLR5. Abolition of TLR5 recognition required extensive boiling time of the bacteria or proteinase K treatment, showing an unusual high stability of the leptospiral flagellins. Interestingly, using antimicrobial peptides to destabilize live leptospires, we detected some TLR5 activity, suggesting that TLR5 could participate in the fight against leptospires in humans or cattle. Using different *Leptospira* strains with mutations in flagellin proteins, we further showed that neither FlaAs nor Fcps participated in the recognition by TLR5, suggesting a role for the FlaBs. These have structural homology to *Salmonella* FliC, and conserved residues important for TLR5 activation, as shown by *in silico* analyses. Accordingly, we found that leptospires regulate the expression of FlaB mRNA according to the growth phase *in vitro*, and that infection with *L. interrogans* in hamsters and in mice downregulated the expression of the FlaBs but not the FlaAs subunits.

Altogether, in contrast to different bacteria that modify their flagellin sequences to escape TLR5 recognition, our study suggests that the peculiar central localization and stability of the FlaB monomers in the periplasmic endoflagella, associated with the downregulation of FlaB subunits in hosts, constitute an efficient strategy of leptospires to escape TLR5 recognition and the immune response.

## Introduction

Leptospires are spirochetal bacteria responsible for leptospirosis, a neglected re-emerging zoonosis [1]. Among the *Leptospira* genus, which includes more than 60 species and 300 different serovars, *Leptospira interrogans* gathers the most pathogenic strains [2]. Rodents and other animals can carry leptospires asymptomatically in the lumen of proximal renal tubules, excrete the bacteria in their urine and contaminate the environment. Vertebrates get infected through skin or mucosa. In most cases, humans have no symptoms or suffer from a flu-like mild disease, but may also show acute severe, potentially fatal, leptospirosis. Antibiotic treatments are efficient only if administered at the onset of symptoms. The high number of leptospiral serovars and strains complicates the diagnosis and impairs vaccinal strategies.

*Leptospira* are motile bacteria able to swim very fast in viscous environments. They possess two endoflagella, one inserted at each pole of the bacteria, which do not protrude outside of the bacteria but are localized and rotate within the periplasmic space. As seen in other spirochetes, the leptospiral genomes exhibit an atypical high number of structural flagellar genes, including four FlaB subunits with homology to FliC, the unique flagellin monomer forming the filament of *Salmonella spp.* The structure of the leptospiral filament and the roles of the different flagellar proteins and additional specific components of leptospires such as the Fcp proteins [3; 4; 5; 6; 7], have been recently elucidated by high-resolution cryo-electron microscopy coupled to model building and crystallography analyses [8]. The leptospiral filament has an atypical flattened helical shape. The four FlaB subunits constitute the core of the flagellum, surrounded by two FlaA and two Fcp subunits that form a sheath. [8].

The innate defense of the host relies on the complement system and on immune receptors, also known as pattern recognition receptors (PRRs), such as Toll-like Receptor (TLR) and NOD-like receptor (NLR) families. TLRs and NLRs recognize conserved microbe-associated molecular patterns (MAMPs) and induce immune inflammatory responses that trigger cellular recruitment, ultimately leading to the destruction of microbes by phagocytes [9].

FliC, the prototypical bacterial flagellin, forms a hairpin-like structure with 4 connected domains designated D0, D1, D2, and D3, with both C and N termini associated in the D0 domain [10]. The D2 and D3 domains are highly variable and support the antigenic diversity. FliC is recognized by different PRRs, expressed on the surface of cells a well as intracellular. TLR5 is expressed at the surface of cells and recognizes monomers of flagellin in the D1 domain, whereas in the cytosol FliC is recognized through the D0 domain by the NAIP inflammasome, which associates with the IPAF/NLRC4, a NOD-like receptor [11; 12]. TLR5 is an essential innate immune receptor expressed in the kidney and, along with TLR4, important to control *Enterobacteria* [13]. Moreover it is one of the very few TLRs able to recognize a protein agonist, conferring potent adjuvant properties, and helping adaptive immune responses [14].

We previously showed that *Leptospira* infection triggers the NLRP3 inflammasome, using the ASC adaptor. The results using ASCKO mice reproduced the results obtained with the NLRP3KO mice and suggest that the contribution of other inflammasome receptors, such as the NAIP/NLRC4 would be minimal [15]. We also showed that *L. interrogans* escapes recognition by human TLR4 [16] as well as murine and human NOD1 and NOD2 [17]. In this work, we investigated whether leptospiral flagellins are either recognized by or also escape recognition by TLR5. Our results suggest that live pathogenic leptospires largely escape recognition by human and murine TLR5, although their FlaBs subunits are able to signal through human TLR5. This suggests that the periplasmic localization of the flagella and the concealing of FlaBs in the core of the filament contribute to avoiding the TLR5 recognition pathway.

## Materials and Methods

### Leptospiral strains and culture conditions

Pathogenic *L. interrogans* serovar Icterohaemorrhagiae strain Verdun, *L. interrogans* serovar Copenhageni strain Fiocruz L1-130, *L. interrogans* serovar Manilae strain L495, and the saprophytic *L. biflexa* serovar Patoc strain Patoc I have initially been provided by the collection of the National Reference Center for Leptospirosis of the Institut Pasteur in Paris. The L495 derivative bioluminescent strain MFLum1 [18], the clinical isolate Fiocruz LV2756 and its non-mobile *fcpA* mutant [5], the *L. interrogans* Manilae *flaA2* mutant, as well as the *flaB4* mutant of *L. biflexa* Patoc have all been previously described [3; 19]. The *L. biflexa fcpA* and *L. interrogans* Manilae *flaB1* mutants have been generated for this study by random mutagenesis [20].

Bacteria were grown in Ellinghausen-McCullough-Johnson-Harris (EMJH) medium (Bio-Rad) at 30 °C without agitation and weekly passaged, counted using a Petroff-Hauser chamber and seeded at 5.10^6^ bacteria per mL (bact/mL). Bacteria in mid-log exponential phase (around 10^8^ bact /mL), and bacteria in stationary phase (around 1 to 5×10^9^ bact/mL) were harvested from 3-6-day old cultures and 10-14-day old cultures, respectively. Unless otherwise specified, experiments were performed with one-week old cultures. The *L. biflexa* Patoc Patoc I strain was passaged twice a week by a 1/250 dilution and seeded at around 5.10^6^ bact/mL. For experiments conducted in New Caledonia, virulent *L. interrogans* serovar Icterohaemorrhagiae strain Verdun was cultured in EMJH medium at 30°C under aerobic conditions as previously described [21]. For *in vitro flaB* gene expression assays, cultures of each *Leptospira* strain were seeded in triplicate at 5.10^6^ (Day 0). On Day 3 (exponential growth phase) and Day 14 (stationary growth phase), 5.10^8^ bacteria from each culture were harvested and centrifuged at 3250 g for 20 min, EMJH was discarded and bacteria were resuspended in 500 μL of RNAlater Buffer (Qiagen) for RNA stabilization, kept at room temperature for 2 h before conservation at −20°C until RNA extraction.

### *In vivo* infection experiments using leptospires

Male and female C57BL/6J mice (7- to 10-week old) were used in this study and were obtained from Janvier Labs (Le Genest, France). TLR5 deficient mice (TLR5KO) in a C57BL/6J background were bred at the Institut Pasteur Paris animal facility and were previously described [18]. Outbred OF1 mice (*Mus musculus*) and golden Syrian hamsters (*Mesocricetus auratus*), initially obtained from Charles River Laboratories, were bred in the animal facility in Institut Pasteur in New Caledonia.

Infections of C57BL/6J mice with *L. interrogans* strains were conducted as described [22]. Just before infection, bacteria were centrifuged at room temperature for 25 min at 3250 ×g, resuspended in endotoxin-free PBS. Leptospires in 200 μL of PBS were injected via the intraperitoneal route (IP) into mice. Experiments were done with sublethal doses of pathogenic *L. interrogans*. Animals were bled at the facial vein sinus (around 50-100 μl of blood, recovered in tubes coated with 20 μl of EDTA 100 mM). A drop of urine was retrieved upon first handling of mice. Animals were killed by cervical dislocation and organs frozen in liquid nitrogen before storage at −80°C or fixed in formaldehyde for histopathology.

Virulence of *L. interrogans* Icterohaemorrhagiae strain Verdun was maintained by cyclic passages in golden Syrian hamsters after intraperitoneal (IP) injection of the LD_100_ at 2×10^8^ leptospires before re-isolation from blood by cardiac puncture at 4.5 days post infection, after euthanasia with CO_2_.

For *in vivo* study of *flaA* and *flaB* gene expression, 6-to 8-week old healthy animals (n ≥ 5 individuals per condition) were infected and experiments were carried out as previously described [21]. Briefly, OF1 mice and hamsters were IP injected with 2×10^8^ virulent *L. interrogans* Icterohaemorrhaghiae strain Verdun in 500 to 800 μL of EMJH medium. After euthanasia with CO_2_, whole blood was rapidly collected by cardiac puncture at 24 h p.i. and conserved in PAXgene blood RNA tubes (PreAnalytiX, Qiagen) for 2 h at room temperature to allow stabilization of total RNA before storage at −20°C until RNA extraction.

### Ethics statement

Animal manipulations were conducted according to the guidelines of the Animal Care following the EU Directive 2010/63 EU. All protocols were reviewed and approved (#2013-0034, and #HA-0036) by the Institut Pasteur ethic committee (CETEA #89) (Paris, France), the competent authority, for compliance with the French and European regulations on Animal Welfare and with Public Health Service recommendations.

### Histology and immunohistochemistry

Transversal sections of kidneys were collected and fixed in formaldehyde 4% for at least 48h at room temperature, embedded in paraffin, and 5 μm thick sections were stained with Hematoxylin-Eosin. Immunohistochemistry was performed on dewaxed sections as described [18]. A rabbit polyclonal serum against the LipL21 (kindly provided by David Haake) was used (1/1000^e^). A Periodic Acid-Schiff (PAS) staining was also associated to the Lip21 immunolabeling to visualize the membranes and brush borders typical of proximal tubules.

### qPCR quantification of leptospiral DNA in blood, urine and organs

The leptospiral load in blood, urine and organs was determined by quantitative real-time PCR (qPCR), as described [22]. Total DNA from blood and urine (around 50 μL) was extracted using a Maxwell 16 automat with the Maxwell blood DNA and cell LEV DNA purification kits (Promega), respectively. DNA was extracted with the QIAamp DNA kit (Qiagen) from organs mechanically disrupted for 3 min at 4°C with metal beads using an automat (Labomodern). Primers and probe designed in the *lpxA* gene of *L. interrogans* strain Fiocruz L1-130 [4] were used to specifically detect pathogenic *Leptospira sp.* [22], using the *nidogen* gene for normalization in kidneys. qPCR reactions were run on a Step one Plus real-time PCR apparatus using the absolute quantification program (Applied Biosystems), with the following conditions for FAM-TAMRA probes: 50 °C for 2 min, 95 °C for 10 min, followed by 40 cycles with denaturation at 95 °C for 15 s and annealing temperature 60 °C for 1 min.

### Reverse and Real-time transcription PCR for cytokine gene expression

Total RNA was extracted from kidneys using the RNeasy mini kit (Qiagen) and RT-qPCR were performed as described [18]. The sequences of primers and probes for IL10, RANTES, and IFNγ have already been described [10][15]. Data were analyzed according to the method of relative gene expression using the comparative cycle threshold (Ct) method also referred to as the 2^**(−ΔΔCt)**^ method. PCR data were reported as the relative increase in mRNA transcripts versus that found in kidneys from naive WT mice, corrected by the respective levels of Hypoxanthine phosphoribosyltransferase (HPRT) mRNA used as an internal standard.

### Total RNA extraction and cDNA synthesis for leptospiral *fla* genes

Total RNA from blood was extracted using a PAXgene blood RNA system from PreAnalytiX (Qiagen). Total RNA from virulent *Leptospira* (4×10^8^ bacteria) cultured *in vitro* at 30 °C and at 37 °C in EMJH medium was also extracted using a High Pure RNA Isolation kit (Roche Applied Science) following the manufacturer’s recommendations. Total RNA samples were treated with DNase (Turbo DNA-Free kit; Ambion, Applied Biosystems) for elimination of residual genomic DNA. Before storage at −80°C, purified RNA was quantified by measurement of the optical density at 260 nm (OD_260_) using a NanoDrop 2000 spectrophotometer (Thermo Fisher Scientific), and the quality of nucleic acids was verified by measurement of the OD_260_/OD_280_ ratio. Then, 1 μg of total RNA was reverse transcribed using a Transcriptor First Strand cDNA synthesis kit (Roche Applied Science) and the provided random hexamer primers for the mix preparation, on a GeneAmp PCR system 9700 instrument (Applied Biosystems) with the following program: 10 min at 25°C; 30 min at 55°C; and 5 min at 85°C. The cDNA synthesized was conserved at −20°C until quantitative PCR (qPCR) assays.

### Quantitative PCR and FlaA and FlaB expression analysis

After cDNA synthesis, qPCR assays were performed using primers purchased from Eurogentec (Seraing, Belgium; Table 1) and specific for the gene coding for the *flaA* and *flaB* subunit genes. Primers were designed using LightCycler Probe Design Software (version 2.0; Roche Applied Science) or the free online Primer3 software (version 0.4.0) using available sequences retrieved from GenBank (NCBI). Amplifications were carried out on a LightCycler 480 II instrument using LightCycler 480 software (v. 1.5.0) and a LightCycler 480 SYBR green I master kit (Roche Applied Science) according to the provided instructions. The amplification program was as follow: a first hot start (95°C for 10 min) and 50 cycles of an activation step at 95°C for 5 s, an annealing step at 62°C for 5 s, and an elongation step at 72°C for 8 s. Each sample was run in duplicate. A single acquisition of fluorescence for calculation of the Ct was processed during the elongation step. The specificity of amplification was verified by size visualization of the PCR product (Table 1) after electrophoresis on a 1.8% agarose gel (Sigma-Aldrich) in 1% TBE (Tris-borate-EDTA) for 30 to 45 min at 120 V and by analysis of the melting curves of the PCR products (melting temperatures, T*m*, in Table 1). All Ct values were analyzed using the qbase^PLUS^ software (Biogazelle, Belgium).

**TABLE 1.**
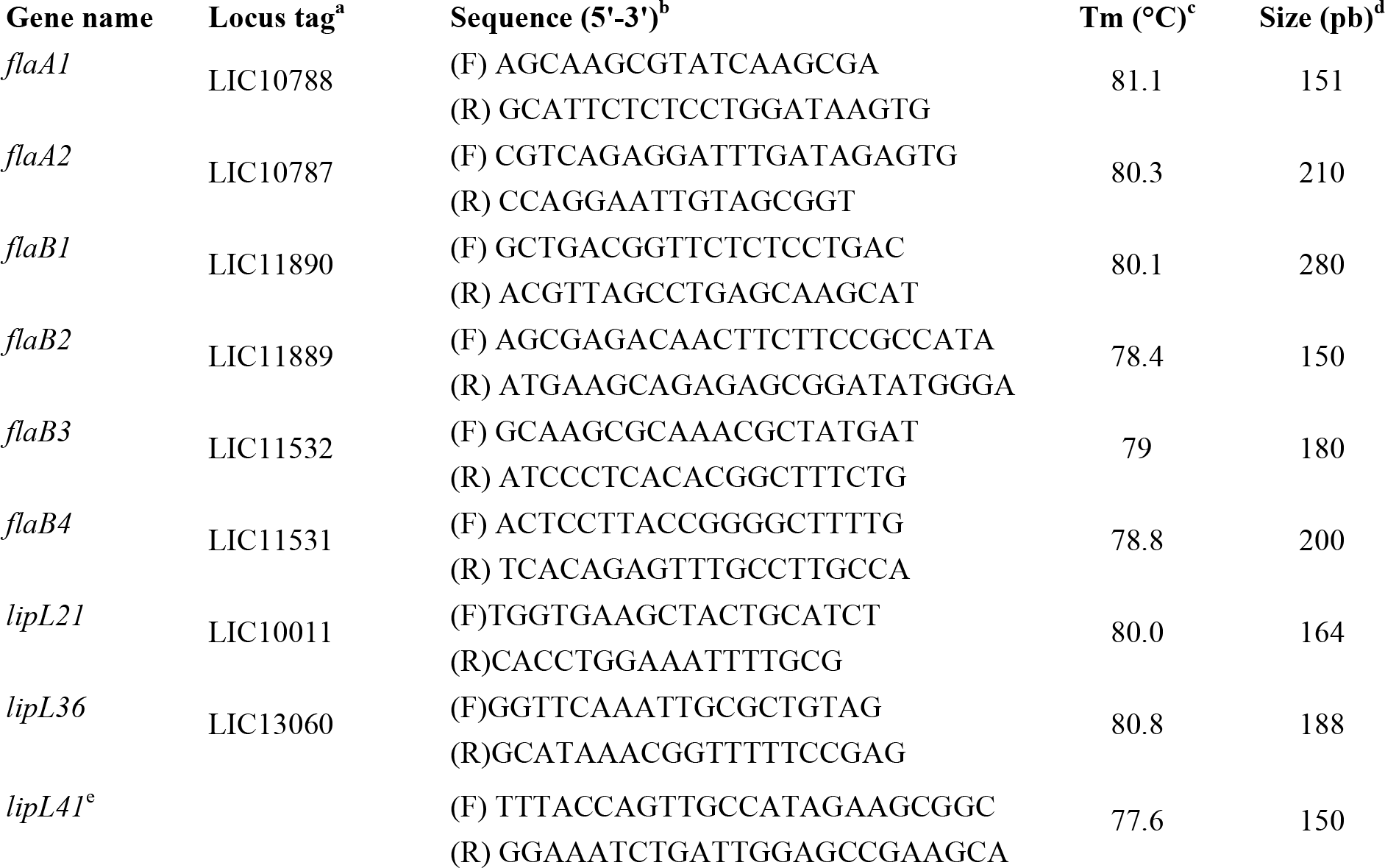
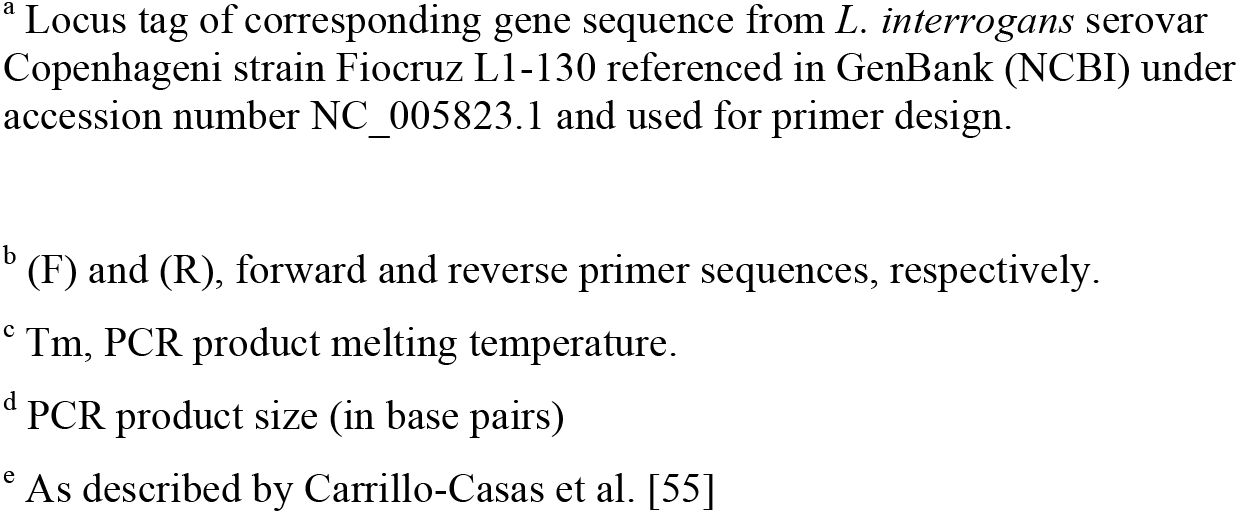
Details and sequence of primers used for qPCR assays.

For *in vivo* infections, the level of expression of each target gene was normalized to the levels of *lipL21*, *lipL36*, and *lipL41* gene, previously validated as reference genes in our conditions [23]. The relative normalized expression ratio was then calculated as the ratio of the *in vivo* to the *in vitro* expression level of bacteria cultured at 30°C. For the *in vitro* bacterial cultures, level of expression of *flaB* genes was normalized to the level of *lipL41* housekeeping gene (Normalized relative quantities).

### Generation of Bone Marrow-derived Macrophage (BMM)

Bone marrow cells (BMC) were obtained as recently described [22]. Briefly, mice were euthanized, femurs dissected, cleaned, and the heads were cut off. BMC were flushed out using a 21G needle to inject culture medium through the bones. BMC were centrifuged (300 g, 5 minutes) and treated with Red Blood Cell Lysis Buffer (Sigma-Aldrich) for 10 minutes, followed by PBS washing. BMC were counted, and 5×10^6^ cells seeded in 100-cm^2^ cell culture dishes in 12 mL RPMI supplemented with 10% fetal calf serum (Lonza), 1X non-essential amino acids (NEEA, Gibco), 1 mM sodium pyruvate (complete medium) supplemented with 1X Antibiotic/Anti-mycotic solution (Gibco) and 10% L929 cell supernatant to provide a source of M-CSF1. Cells were incubated for 7 days at 37°C with 5% CO_2_. At day 3, 5 mL of the same medium was added. At day 7, the medium was removed, and 3 mL of cell dissociation buffer (Gibco) was added to harvest the bone marrow macrophages (BMMs). BMMs were collected by scrapping, centrifuged, enumerated and seeded in 96-well plates at a density of 2×10^5^ cells per well in complete medium without antibiotics. BMMs were rested for 2 to 4 h and stimulated for 24 h with different leptospiral strains, live or heat-killed for 30 min at 100 °C, at a MOI of 1:100, or 1:50 or with 100 ng/mL of controls [Standard Flagellin from *Salmonella typhimurium* (FLA) and LPS *E. coli* ultra-purified (both from InvivoGen)]. The keratinocyte-derived (KC/CXCL1) cytokine was measured in cell supernatants 24 h post-stimulation, by ELISA using Duo-Set kits (R&D Systems), according to the supplier’s instructions.

### TLR5/ NF-κB assay in human epithelial cell line HEK-blue-KD-TLR5

Human embryonic kidney TLR5 knock-down cells (HEK-Blue-KD-TLR5 cells, Invivogen) were used for the *in vitro* experiments In these HEK-BLUE cells, the activation of NF-κB drives the expression of the alkaline phosphatase enzyme that induces a color shift from pink to blue of the chromogenic substrate in the HEK-Blue Detection Media (Invivogen). These cells were cultured in complete DMEM medium composed of DMEM GlutaMAX (Gibco) with 1 mM sodium pyruvate (Gibco), 1X NEEA (Gibco) and 10 % V/V heat-inactivated foetal calf serum (Hi FCS, Gibco). On day 1, cells were detached by 1 min incubation in cell dissociation buffer (Gibco) followed by gentle flush with medium. Cells were then seeded in 22.1 cm^2^ cell culture dishes (TPP) at less than 30 % confluence and incubated overnight at 37 °C, 5 % C0_2_. Cell transfections were performed on day 2, whilst the cells remained under 60 % confluence and with a total amount of 3 μg of DNA per dish. For each dish, between 100 ng to 1 μg of pUNO1-humanTLR5, pUNO1-murineTLR5 (Invivogen), pcDNA3.1-bovine TLR5 [24] or the corresponding empty vector was used, complemented up to 3 μg with pcDNA3.1. The transfection reagent 1X FuGENE HD (Promega) in serum free OptiMEM (Gibco) was incubated during 25 min with the DNA followed by transfection of the cells according to the manufacturer’s instruction. On day 3, transfected HEK-Blue-KD-TLR5 cells were stimulated in 96-wells plates. Briefly, 20 μL Flagellin from *Salmonella typhimurium* as a control (Standard FLA-ST (Invivogen) or leptospires resuspended in PBS at a MOI between 1:50 to 1:200 were added in empty wells. Transfected HEK-Blue-KD-TLR5 cells were then gently flushed in PBS and resuspended in HEK-Blue Detection Media (Invivogen) at 2.8 × 10^5^ cells/mL. 180 μL of cell suspension, corresponding to 50 000 cells, were then added on top of the agonists in each well and plates were incubated for 24h at 37°C, 5 % CO_2_. In each well, the activation of NF-κB through TLR5 was assessed by absorbance measurements at 630 nm. All heat treatments were performed under agitation at 300 rpm and in PBS on the diluted leptospires preparations right before addition in the wells. Proteinase K treatments of leptospires (from *Tritirachium album*, Qiagen) were performed under agitation at 300 rpm in PBS for 2h at 37°C, to avoid killing the leptospires. Such treatment was followed by heat inactivation of the enzyme and bacteria at 100°C for 30 min. The non-inactivated fraction and mock treatment without leptospires were also tested on HEK-Blue-KD-TLR5 cells. Treatment of live leptospires was performed with 100 μg/mL of the human cathelicidin LL37 (Invivogen) in PBS for 2 hours.

### *In silico* analyses of the flagellin protein sequences

All the *in silico* analyses were performed using either Uniprot or GeneBank available sequences. All corresponding accession numbers are mentioned in the figure legends. Amino acid sequence homology percentage (identity) was obtained using BLAST. Alignments of the sequences were performed with MEGA X [25] and using the Clustal method. Structural predictions based on amino acid sequences were obtained using the Phyre2 [26] and figures colored and modified with Chimera [27].

### Statistical analyses

Statistical analyses were performed using non-parametric Mann-Whitney test, that does not assume a normal distribution of the samples, and stars were attributed according the following p values: * p<0.05; ** p<0.01.

## Results

### TLR5 deficiency does not modify the course of leptospirosis in mice

To study the potential involvement of the TLR5 receptor in the host defense against leptospires, we used a murine model of leptospirosis and compared the susceptibility of C57BL6/J (WT) mice *versus tlr5* knock-out (TLR5KO) mice in the same genetic background after intraperitoneal infection with a sublethal dose of 10^7^ *L. interrogans* (serovar Manilae strain L495). Leptospiral loads were measured by q-PCR in blood and urine (Figure 1A) and organs (Figure 1B) 3 days, 7 days and 15 days post-infection (p.i). As previously described [18; 28], leptospires were present in blood, liver, spleen, lungs and kidney at day 3 p.i (Figure 1A and 1B). At day 7 p.i, leptospires were detected in urines, but not in blood or kidneys, similar as previously observed [29]. At day 15 p.i, leptospires were present in urine and kidneys. No difference of leptospiral loads could be observed between WT and TLR5KO mice in blood, urine or organs. In addition, mRNA expression of pro-(IFNγ), anti-(IL-10) inflammatory cytokines and RANTES chemokine measured by RT-qPCR at day 3 and 7 p.i in the kidneys did not differ between WT and TLR5KO mice (Figure 1C). Altogether these results suggest that the presence of TLR5 does not play a major role in the murine defense against experimental leptospirosis.

**Figure 1.**
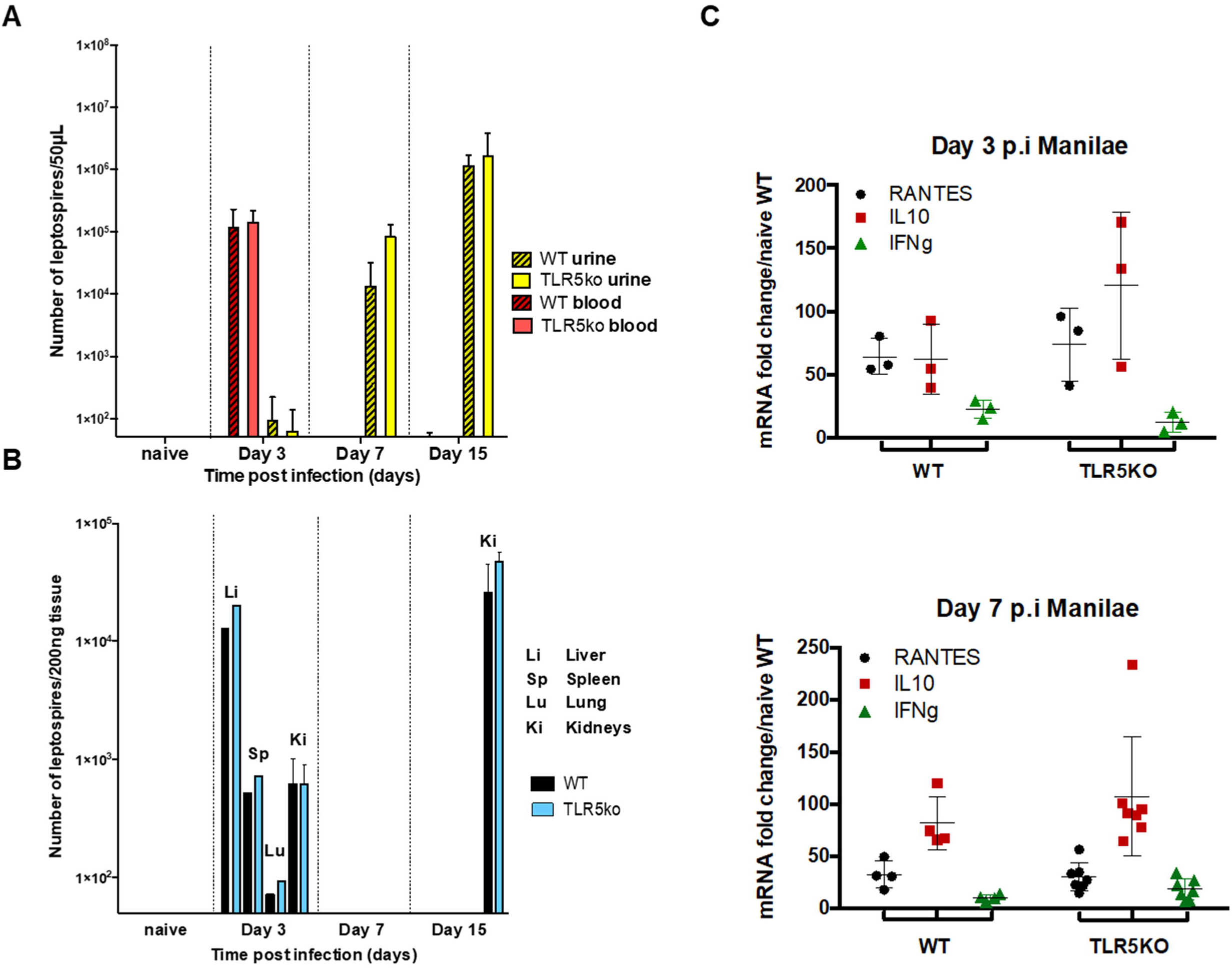
Lack of TLR5 does not modify the course of leptospirosis in mice. Intraperitoneal infection of 7-week old female C57BL/6J (WT) mice (n= 4) and TLR5KO (n= 4) mice with 10^7^ *L. interrogans* Manilae (strain L495). **A, B**) Bacterial loads determined by q-PCR of leptospiral DNA at different days post infection (p.i), **A**) in blood (red) and urine (yellow) in WT (empty bars) and TLR5KO mice (hatched bars); **B**) in organs [liver (Li), spleen (Sp), Lung (Lu) and Kidney (Ki)] from WT (black bars) and TLR5KO (blue bars). **C**) Inflammation measured in kidney by RT-qPCR of cytokines (RANTES, IL10, IFNγ),at 3 days p.i and 7 days p.i.

### TLR5 deficiency does not change the localization of leptospires in kidneys

Because TLR5 is expressed in epithelial renal cells and plays an important role to control bacteria responsible for urinary tract infection [13], we hypothesized that the absence of TLR5 could impact the localization of leptospires in the kidneys or the induced nephritis [18; 28]. We used the bioluminescent derivative MFLum1 of *L. interrogans* Manilae strain L495 to visualize leptospires in the kidneys 15 days post-infection. Similar as described for the bacterial loads measured by q-PCR, levels and shape of emitted light, reflecting live bacteria [29], were equivalent between WT and TLR5KO infected mice (Figure 2A). Using immunohistochemistry, we further investigated the presence of leptospires in kidneys of WT and TLR5KO mice. Minimal inflammatory cellular infiltrates were noted with similar incidence and severity in the cortex of both WT and TLR5KO infected mice (Figure 2B a-f), whereas no inflammation was observed in the naive WT control. Labeling of leptospires with an anti-LipL21 antibody [17] revealed a low number of *Leptospira*-infected tubules in the renal cortex, as already described [18] (Figure 2B g-i). In histological sections of the kidney stained with Periodic Acid-Schiff, we only found leptospires in some proximal tubules, associated with the PAS positive microvilli of the brush border at the luminal surface of the tubular epithelium, as previously described in rats [30]. No differences were observed between WT and TLR5KO mice (Figure 2B j-k). Altogether these results suggest that TLR5 does not play any major role in host protection against *L. interrogans* Manilae L495 infection in mice and that leptospires may therefore escape the TLR5 response.

**Figure 2.**
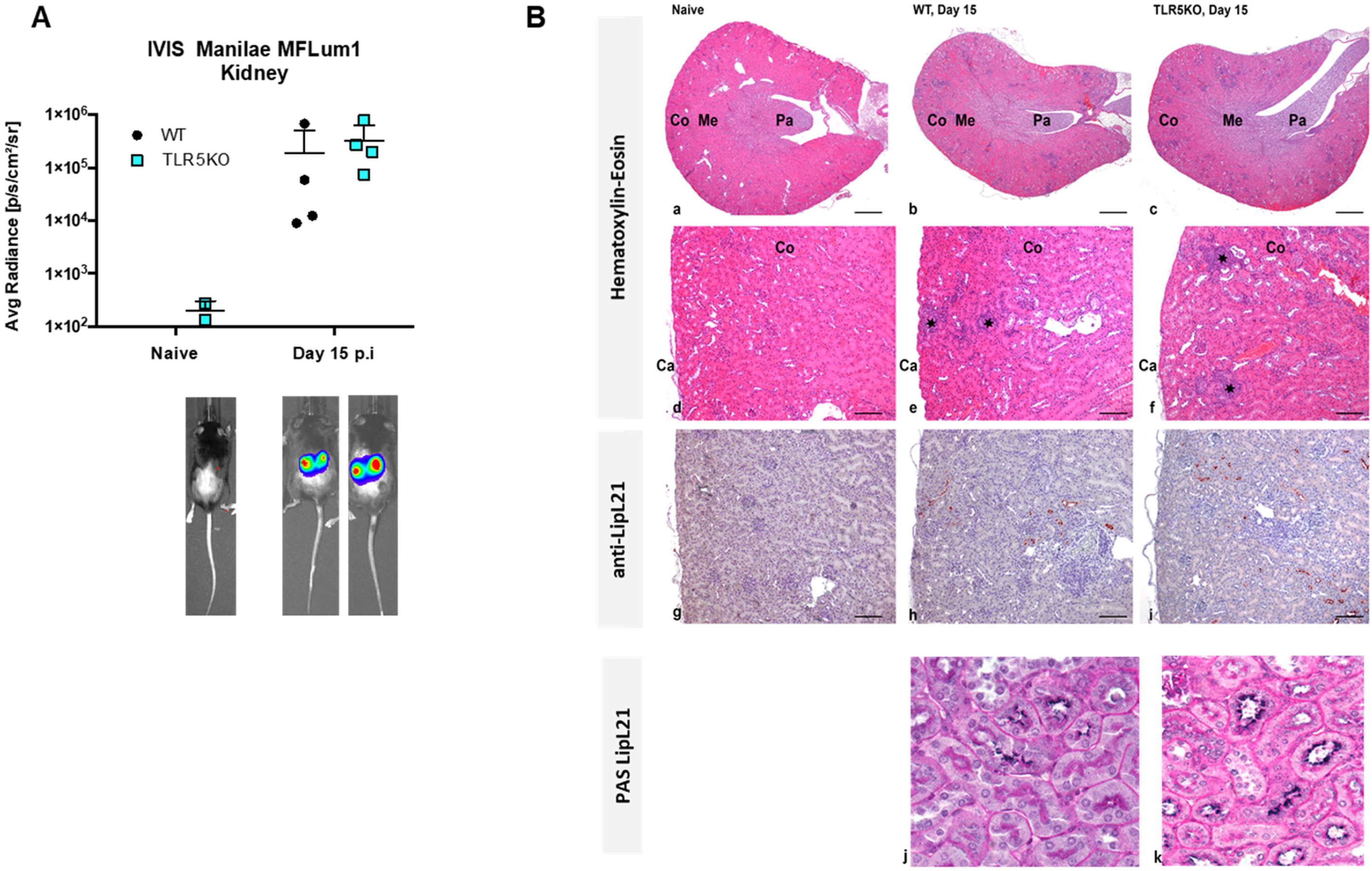
Lack of TLR5 does not modify the kidney colonization. **A)** Tracking by live imaging (IVIS) of 7-week old C57BL/6J mice infected intraperitoneally with 10^7^ *L. interrogans* Manilae (strain L495) bioluminescent derivative MFLum1. The imaging has been performed 15 days post-intraperitonal infection on anesthetized mice after luciferin administration. The graph represents the mean ± SEM of the average radiance of n = 4 mice in each group, imaged in dorsal position, and gated on the whole body. The background level of light was measured on a control TLR5KO mice injected with PBS at the time of infection. **B)** TLR5 deficiency does not modify the localization of leptospires in the proximal tubules. Histological sections and immunolabeling of the kidneys of naive TLR5KO, infected WT and TLR5KO mice 15 days p.i. **a-c)** Kidney, Hematoxylin-Eosin stain, Original magnification x2, Scale bar: 500 μm. Cortex (Co), Medulla (Me), Papilla (Pa), Capsule (Ca). **d-f)** Kidney cortex, Hematoxylin-Eosin stain, Original magnification ×10, Scale bar: 100 μm. The stars indicate the focal inflammatory infiltrates. **g-i)** Anti-LipL21 labelling of leptospires in renal tubules, Original magnification ×10, Scale bar: 100 μm. **j,k)** Double labelling LipL21/Periodic Acid-Schiff (PAS) to stain the PAS positive brush borders present in proximal tubules only. Original magnification x40, Scale bar: 25 μm.

### Live pathogenic leptospires do not signal through murine and human TLR5 *in vitro*

Bone marrow derived macrophages (BMMs) from WT and TLR5KO mice were infected with different live strains of *L. interrogans* and the production of the KC chemokine was measured by ELISA 24h p.i in the cellular supernatants. We did not find any difference between both genotypes (Figure 3A), which correlated with the *in vivo* experiments and indeed strongly supporting the observation that live leptospires were not recognized through the murine TLR5.

**Figure 3.**
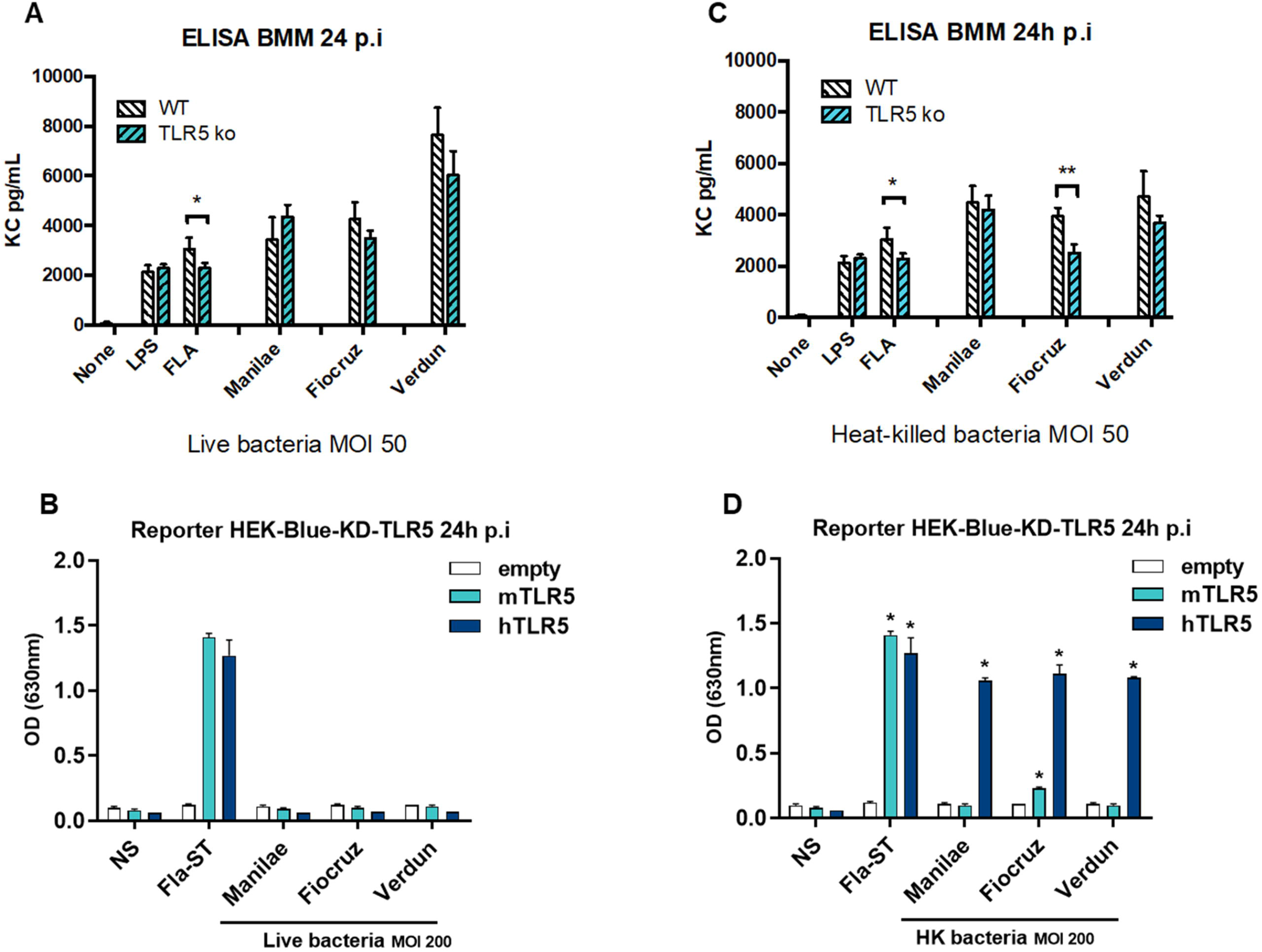
Live pathogenic leptospires do not signal through TLR5, although heat-killed leptospires signal via human TLR5, but barely through murine TLR5. **A,C**) KC production measured by ELISA in the supernatants of BMM from WT (hatched empty) and TLR5KO (hatched blue) mice 24 hours p.i with different serovars of virulent *L. interrogans* at MOI 50 (Manilae strain L495, Copenhageni strain Fiocruz L1-130, Icterohaemorrhagiae strain Verdun. **A**) Live bacteria, **C**) Heat killed bacteria (30 min 100 °C). LPS from *E. coli* (100 ng/mL) and unpurified FLA from *Salmonella typhimurium* (500 ng/mL) were used as controls. Each graph corresponds to one representative experiment out of 3 independent experiments. Data are expressed as the mean (+/-SD) of pooled BMM preparations from n=3 mice with 4 technical replicates, and statistics analyses were performed using non-parametric Mann-Whitney test: p<0,05*; p<0,01 **. **B, D**) Reporter NF-κB assays in HEK-Blue-Knock Down (KD)-TLR5 cells transfected with the human TLR5 (dark blue bars), mouse TLR5 (light blue bars), or empty plasmid (empty bars) and stimulated for 24 h with **B**) live bacteria and **D**) heat-killed bacteria at MOI 200. Flagellin from *Salmonella typhimurium* (Fla-ST) at 500 ng/mL was used as control. Data are expressed as the mean of OD630 for n=3 replicates and represent one out of 3 independent experiments.

We assessed next whether the escape of recognition by murine TLR5 is a species-specific phenomenon. Indeed, we previously highlighted PRR species-specificities of leptospiral MAMPs recognition, such as murine TLR4 receptor that recognizes the leptospiral LPS, whereas human TLR4 does not, and conversely the human NOD1, but not the murine NOD1 is able to sense leptospiral muropeptides [17; 31]. We therefore transfected HEK-Blue-KD-TLR5 cells with either human TLR5, murine TLR5, or an empty control vector. No signal corresponding to murine or human TLR5-mediated NF-κB activation was obtained upon infection with different live *L. interrogans* strains at MOI of 10 and 100 (date not shown) and neither at a high MOI of 200 (Figure 3B), which suggested that leptospires also escape the human TLR5 recognition.

### Heat-killed leptospires signal through human TLR5 but barely through murine TLR5

The specificity of TLR5 activation is usually assessed by inactivating and denaturing a potential ligand through heat-inactivation. Here, we observed that Manilae L495 and Icterohaemorrhagiae Verdun strains inactivated at 100°C for 30 min induced equivalent levels of KC in WT and TLR5KO murine BMMs, which was consistent with the results obtained using live bacteria (Figure 3A). In contrast, the heat-killed Copenhageni Fiocruz strain L1-130 unexpectedly induced less cytokines in TLR5KO than in WT BMM (Figure 3C), suggesting that an agonist present in the inactivated Fiocruz strain could be recognized by murine TLR5. Unexpectedly, heat-killed leptospires from all serovars strongly activated HEK-Blue-KD-TLR5 transfected with human TLR5 (Figure 3D). Further, despite the fact that both Manilae L495 and Icterohaemorrhagiae Verdun strains did not stimulate murine TLR5, a slight activation signal was observed with the Copenhageni Fiocruz L130 strain, which was consistent with the cytokine results in BMMs (Figure 3C). The experiment was performed in parallel with an empty plasmid, showing that these results were indeed specific to TLR5 activation, and did not depend on a different NF-κB activation pathway (Figure 3D). Altogether these unexpected results suggested that only heat-killed leptospires can signal through human TLR5, but not or only barely through murine TLR5, providing a new example of species-specificity of PRR recognition of leptospiral MAMPs.

### A heat-resistant protein from heat-killed leptospires signals through TLR5

To our knowledge, our results showing TLR5 activation using heat-inactivated leptospires has never been described before. Thus, we first ensured that the signal observed was indeed attributed to flagellin-like proteins of leptospires interacting with TLR5. Since only stimulation with heat-killed bacteria resulted in TLR5 signaling, we anticipated that a proteinase K treatment would destroy the protein involved in the signaling. Therefore, we treated live and heat-killed Fiocruz L1-130 leptospires with proteinase K, followed or not by heating at 100 °C for 30 min to inactivate the enzyme. We stimulated TLR5 transfected HEK-Blue-KD-TLR5 with these preparations and a mock control without bacteria. Although the proteinase K treatment had no effect on live bacteria, it decreased the signal on heat-killed bacteria (Figure 4A). In contrast to live bacteria treated with proteinase K and subsequently heated, which resulted in a strong TLR5 activation, TLR5 signaling was not restored in heat-killed bacteria treated with proteinase K, suggesting that the proteinase K had digested all TLR5 agonists (Figure 4A). This experiment confirmed the protein nature of the agonist present in heat-killed leptospires, which was not affected by proteinase K activity in live bacteria. We hypothesize that in live leptospires, the periplasmic location of the endoflagella would protect the flagellin subunits from proteinase K digestion, thus potentially explaining why live bacteria do not signal through TLR5 and are not affected by the enzyme (Figure 4B).

**Figure 4.**
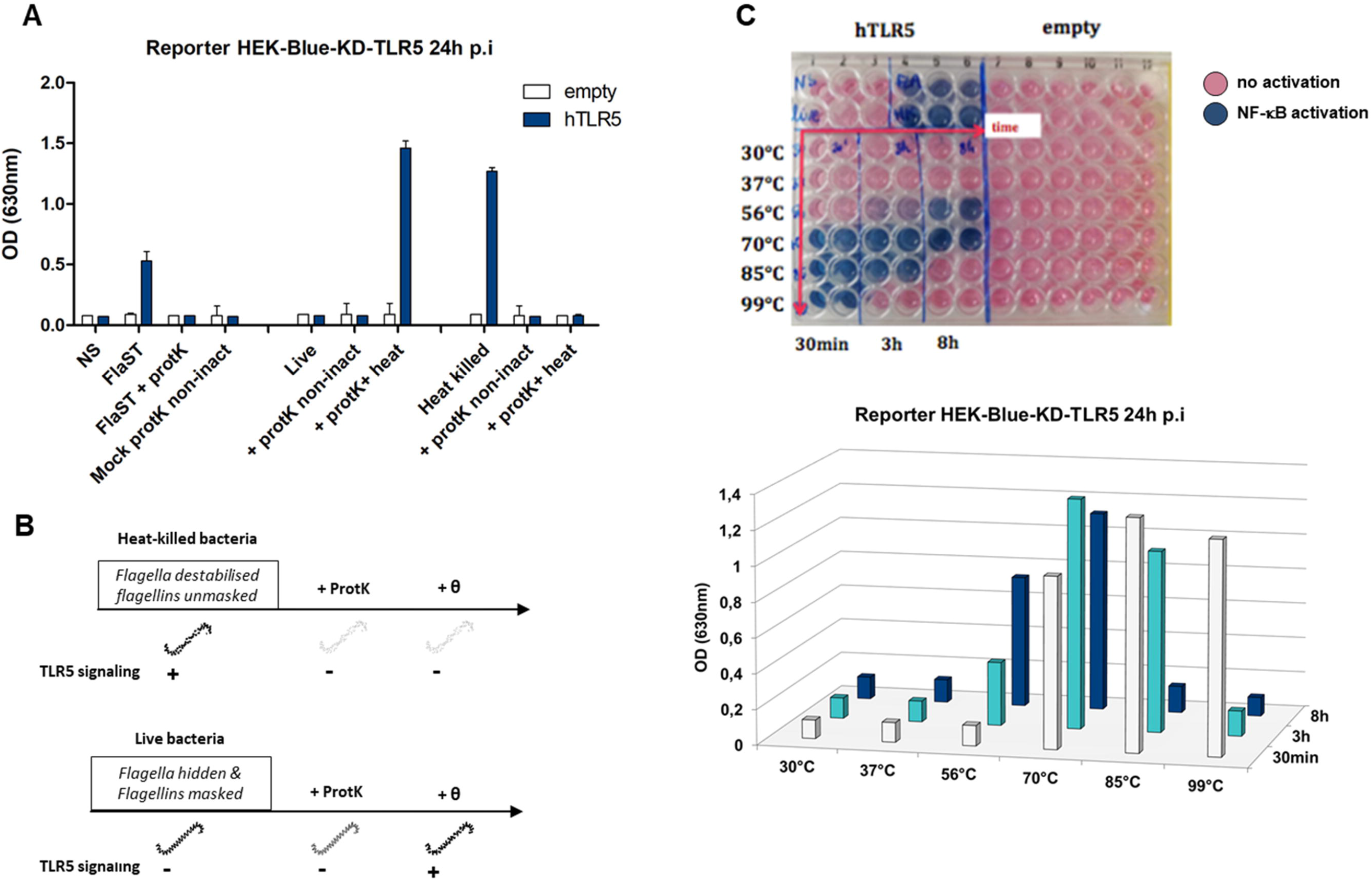
A very stable protein from leptospires signal through TLR5. **A)** NF-κB reporter assay in HEK-Blue-KD-TLR5 cells transfected with the human TLR5 (dark blue bars), or empty plasmid (empty bars) and stimulated with live and heat killed *L. interrogans* Copenhageni strain Fiocruz (MOI 100) treated or not with Proteinase K (protK) followed or not by heat inactivation at 99°C for 30 minutes (inact or non-inact). Flagellin from *Salmonella typhimurium* (Fla-ST) at 500 ng/mL was used as control. Data are expressed as the mean OD630 of n=3 replicates and represent one out of 3 independent experiments. **B)** Chronogram of proteinase K experiments. **C**) Picture of the plate 24 hours p.i. (upper panel) and corresponding graph (lower panel) of Reporter NF-κB assays in HEK-Blue-KD-TLR5 cells transfected with the human TLR5 or empty plasmid and stimulated with *L. interrogans* Copenhageni strain Fiocruz L1-130 at MOI 100 incubated at various temperatures during 30 min (empty bars), 3 hours (light blue bars) or 8 hours (dark blue bars). Data are expressed as the mean of OD630 of n=2 replicates and represent one out of 3 independent experiments.

Next, we investigated the unusual thermostability of the TLR5 agonist, by incubating live Fiocruz L1-130 leptospires at different temperatures (from 30°C and up to 99°C) and for different durations (from 30 min and up to 8 h) (Figure 4C) before stimulation of HEK-Blue-KD-TLR5 transfected with human TLR5. Interestingly, after 8 h incubation at 30°C (the optimal temperature for leptospires’ growth *in vitro*) or at 37 °C (the host temperature), we did not observe any TLR5-dependent signaling. Of note, at 56°C, the usual temperature to inactivate leptospires [15; 18] whilst keeping the leptospiral shape integrity, a signal started after 3 hours of incubation, but even 8 hours were not enough to get a full TLR5 signaling. At 70°C, the temperature classically used to depolymerize the *Salmonella*’s flagellum filament [10], a 30 min incubation was sufficient to stably activate TLR5 for 8 hours. The signal observed with leptospires incubated for 30 min at 85°C disappeared after 8 hours, whereas the positive signal observed after heating the bacteria at 99°C for 30 min disappeared after 3h of heating (Figure 4C). These results confirmed the protein nature of the TLR5 agonist of leptospires, since the activation can be extinguished by heating the bacteria for an extended time at high temperature. Interestingly, these data suggest that, like other bacteria, the leptospiral flagellum depolymerizes at 70°C, which allows for the release of active monomers recognized by the human TLR5. However, in contrast with the *Salmonella* FliC, which is inactivated after 15 min at 100°C, leptospiral flagellins appear to be highly resistant to heating.

### Antimicrobial peptides destabilize live leptospires and unmask a TLR5 signal

Since we revealed the potential for TLR5 recognition of leptospires by heating at high non-physiological temperatures, we wondered whether leptospires could signal through TLR5 after being destabilized or killed with antibiotics or antimicrobial peptides. Antibiotic treatments (at MIC concentrations) including gentamicin, azithromycin and penicillin G, the latter being known to target the cell wall, did not induce any TLR5 signal (data not shown). Next, we used cathelicidin LL37, an antimicrobial peptide which has been shown to be active against leptospires [32], and that was recently associated with a better outcome in human patients with leptospirosis [33]. Furthermore, LL37 has also been shown to prevent death in young hamsters experimentally infected with the Fiocruz L1-130 strain [33]. In our study, live *L. interrogans* Manilae L495 bacteria pre-treated with LL37 induced a modest but significant TLR5 signal (Figure 5A). Another antimicrobial peptide, the bovine Bmap28 has been described to be 50 to 100 times more efficient in killing leptospires than LL37 [32]. We thus tested whether bovine TLR5 could recognize leptospires. Interestingly, although as already published [24], bovine TLR5 did not recognize the *Salmonella* flagellin that we used as a positive control of activation, we found that it recognized heat-killed Fiocruz L1-130 (Figure 5B). The magnitude of the bovine TLR5 signaling was intermediate between the weak response observed with murine TLR5 and the response seen when using human TLR5 (Figure 5B). However, rather than reflect real differences between bovine and human TLR5, this lower response may actually result from the heterologous expression system of bovine TLR5 in the human HEK cell system, that has been shown to impact the responsiveness [34]. Therefore, we anticipate that Bmap28 could allow for a TLR5 signaling in bovines infected with *Leptospira* spp.

**Figure 5.**
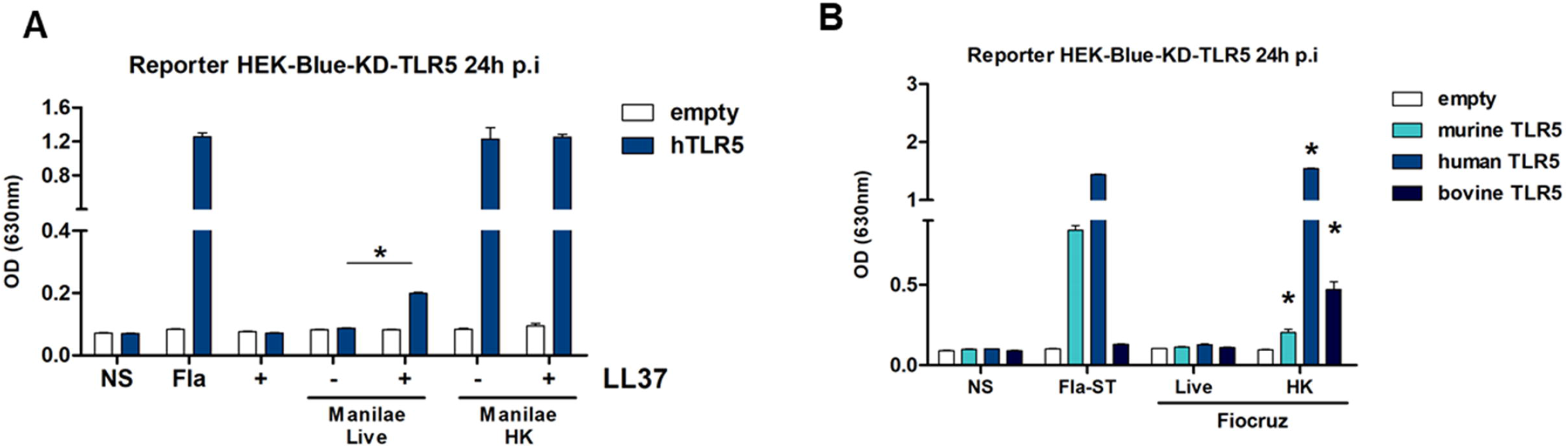
Antimicrobial peptides unmask the leptospiral ability to signal through TLR5, and bovine TLR5 recognizes leptospires. **A)** Reporter NF-κB assay in HEK-Blue-KD-TLR5 cells transfected with the human TLR5 (dark blue bars), or empty plasmid (empty bars) and stimulated with live and heat killed *L. interrogans* strain Manilae (MOI 100) treated or not with human cathelicidin (LL37) at 100 μg/mL for two hours before stimulation. **B)** Reporter NF-κB assay in HEK-Blue-KD-TLR5 cells transfected with the mouse TLR5 (light blue bars), human TLR5 (dark blue bars), bovine TLR5 (black bars) or empty plasmid (empty bars) and stimulated with live and heat killed *L. interrogans* Copenhageni (strain Fiocruz L1-130). Flagellin from *Salmonella typhimurium* (Fla-ST) at 500 ng/mL was used as control. Data are expressed as the mean OD630 of n=3 replicates and represent one out of 3 independent experiments and statistics analyses were performed using non-parametric Mann-Whitney test: p<0,05*.

### *In silico* analyses of potential TLR5 binding sites within leptospiral FlaBs

Two leptospiral *flaA* (*flaA1* and *flaA2*) and four *flaB* genes (*flaB1* to *flaB4*) have been annotated in the *L. interrogans* genomes according to their similarity with the *Salmonella* flagellin (FliC), the two families sharing respectively around 25 % and 38 % identity at the protein level with FliC (Figure 6A). Structural studies recently showed that the FlaBs subunits constitute the core of the flagellum, and the other subunits constitute an asymmetric outer sheath, with FlaBs interacting with FlaAs on the concave site and with FcpA on the other side of the curvature. FcpA and FcpB associate in a lattice forming the convex part of the endoflagellum [8] (Sup Figure 2A). Using the BLAST-P software, we found that the different FlaB subunits from the *L. interrogans* Fiocruz strain share 57% to 72% of identity and most probably result from gene duplication events (Sup Figure 1A). Similar results were obtained with the saprophytic *L. biflexa* Patoc strain (Sup Figure 1A). We then used the Phyre2 software to model the FlaBs structures according to their primary amino acid sequences, using the FliC protein sequence as a base. FliC folds in four regions D0 to D3, forming an inverted L shape (Figure 6B), with both N-term and C-term in the D0 domain. Region D1, in the inner face of the monomer, is involved in the interaction of FliC with the leucine–rich repeat (LRR) domains of TLR5 via 3 binding sites (Figure 6A, 6B and Sup Figure 2B)[35; 36]. There is also a region in the C-term part of the D0 domain that is not directly involved in the binding to TLR5 but important for the stabilization of the TLR5 dimers upon binding to FliC (Figure 6A, 6B and Sup Figure 2B) [37]. All FlaB subunits from *L. interrogans* and *L. biflexa* harbor orthologues of the D0 and D1 domains of FliC, while missing the D2 and D3 domains (Figure 6C, and data not shown). We also checked whether FlaA1 or FlaA2 could have a structure mimicking the D2-D3 domains of FliC, but leptospiral FlaA1 and FlaA2 looked globular, mainly presenting ß sheets and do not resemble the missing domains (Figure 6C). Interestingly, we found that the FlaBs possessed the 3 conserved sequences important for TLR5 binding in the D1 domain (Figure 6C and 6D). Then, we compared the different pathogenic *L. interrogans* and the saprophytic *L. biflexa* Patoc I strain and found that the four FlaBs, although distinct from each other (Sup Figure 1A), were highly conserved in the consensus regions of the TLR5 binding domains in D1 (99 to 100% identity among the different pathogenic serovars, the Patoc FlaBs being less conserved (Figure 6D)). We also found in FlaBs the consensus in the D0 domain involved in the flagellin /TLR5 complex stabilization (Figure 6C). We compared the leptospiral FlaB sequences in these 3 consensus binding TLR5 regions with other spirochetes, *Borrelia burgdorferi* and *Treponema* spp, the latter known to signal via TLR5 when FlaBs are expressed as recombinant proteins [38] and also with bacteria known to dodge the TLR5 response such as *Helicobacter pylori* [39] and *Bartonella bacilliformis* [10], presenting variations in those consensus sequences of their flagellins (Sup Figure 3A). In addition, we also found this FlaB region to be 100 % conserved in a panel of major species of *Leptospira* circulating all over the world, including potential human pathogens, such as *L.borgpeterseni, L. kirschneri*, *L. noguchii*, *L. weilii*, *L. santarosai*, as well as *L. licerasiae*, belonging to another clade of species of lower virulence [2](Sup Figure 3B). These alignments show that the TLR5 binding site region is highly conserved in all leptospiral FlaBs. Therefore each of the four FlaB subunit could potentially signal through TLR5, since leptospiral FlaBs share the 2 first consensus with TLR5 activating bacteria and the different residue observed in the consensus number 3 is also present in TLR5-activating *Treponema* flagellins [40].

**Figure 6.**
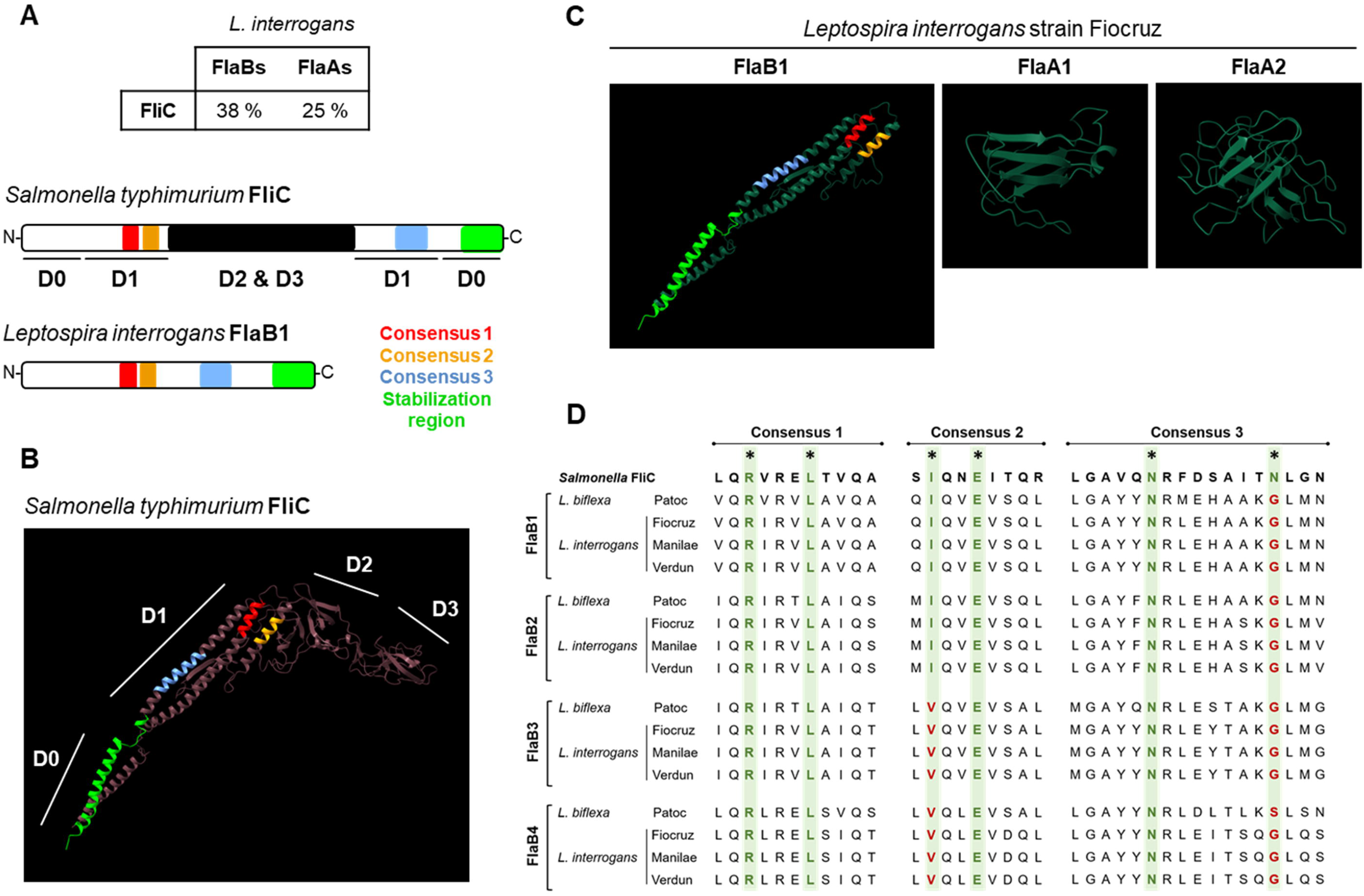
Comparison of leptospiral Flagellins and FliC structures in relation with TLR5. **A)** Amino acid sequence homology average percentage between *Salmonella typhimurium* FliC (P06179) and *Leptospira interrogans* Copenhageni (strain Fiocruz L1-130) FlaBs (LIC11531, LIC11890, LIC11889 and LIC 11542) and FlaAs (LIC10788 and LIC10787) and primary structures of the flagellin proteins with TLR5 binding consensus. **B)** *In silico* (Phyre2 and Chimera softwares) prediction of *Salmonella typhimurium* FliC (P06179) structure with the four described domains and with positions of the TLR5 binding consensus: 1 (red), 2 (yellow) and 3 (light blue) and stabilization region (light green) highlighted. **C)** *In silico* (Phyre2 and Chimera softwares) prediction of *L. interrogans* Copenhageni (strain Fiocruz L1-130) FlaB1 (LIC11531) with the positions of the TLR5 binding consensus and stabilization region highlighted, FlaA1 (LIC10788), FlaA2 (LIC10787). **D)** Clustal (MEGA software) alignment of the amino acid sequences for the TLR5 binding consensus regions of: *Salmonella enterica* FliC (GeneBank QDQ31983.1), *L. biflexa* Patoc (strain Patoc I) FlaB1 (LEPBIa1589), FlaB2 (LEPBIa2133), FlaB3 (LEPBIa2132), FlaB4 (LEPBIa1872), *L. interrogans* Copenhageni (strain Fiocruz L1-130) FlaB1 (LIC11531), FlaB2 (LIC18890), FlaB3 (LIC11889), FlaB4 (LIC11532), *L. interrogans* Manilae (strain L495) FlaB1 (LMANv2_590023), FlaB2 (LMANv2_260016), FlaB3 (LMANv2_260015), FlaB4 (LMANv2_590024) and *L. interrogans* Icterohaemorrhagiae (strain Verdun) FlaB1 (AKWP_v1_110067), FlaB2 (AKWP_v1_110429), FlaB3 (AKWP_v1_110428) and FlaB4 (AKWP_v1_110068).

### FlaBs, but not FlAs or Fcps induce TLR5 signaling

To confirm the putative role of the FlaBs subunits in inducing TLR5 signaling, we used different mutants deficient either in FlaAs, FlaBs or Fcps subunits to stimulate HEK-blue reporter cells transfected with human TLR5. Of note, both *flaAs* and *fcps* genes are in operons, and the *flaA2* mutant lacks both FlaA1 and FlaA2 subunits [3]. Likewise, the *fcpA* mutant lacks both FcpA and FcpB subunits [5; 6]. Our results showed that the TLR5 signaling induced with the heat-killed *fcpA* mutants in Patoc I or in Fiocruz L1-130 was equivalent to the activation observed with parental strains (Figure 7A and 7B). TLR5 signaling was not changed with the *flaA2* mutant in Manilae L495, however the *flaB1* mutant induced a lower activation (Figure 7C). We also observed a decrease of the TLR5 response with the Patoc I *flaB4* mutant (Figure 7D). These results suggest that the FlaB subunits, but not the FlaAs or Fcps, are involved in the TLR5 signaling, and are in line with sequence comparison data.

**Figure 7.**
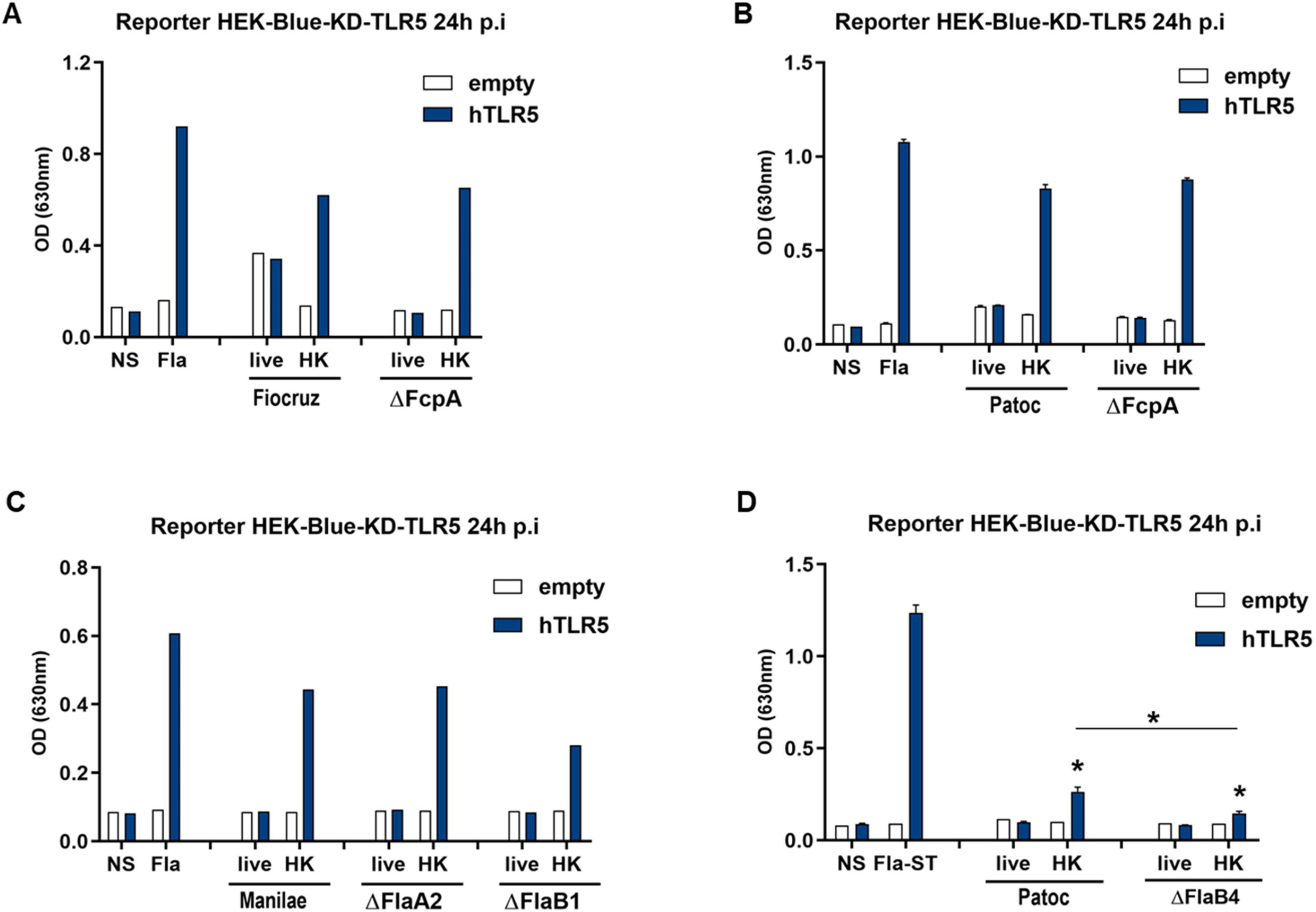
FlaB flagellins are responsible for the signaling, not FlaAs nor Fcps. **A-D)** NF-κB reporter assay in HEK-Blue-KD-TLR5 cells transfected with the human TLR5 (dark blue bars), or empty plasmid (empty bars) and stimulated at a MOI 100 with live or heat killed **A)** *L. interrogans* strain Fiocruz LV2756 or the ΔFcpA strain, **B)** *L. biflexa* strain Patoc and ΔFcpA mutant, **C)** *L. interrogans* strain Manilae and ΔFlaA2 or ΔFlaB1 mutants, **D)** *L. biflexa* strain Patoc and ΔFlaB4 mutant. Flagellin from *Salmonella typhimurium* (Fla-ST) at 500 ng/mL was used as control. Data are expressed as the mean OD630 of n=2/3 replicates and represent one out of 3 independent experiments. Statistics analyses were performed using non-parametric Mann-Whitney test: p<0,05*, and comparing stimulated cells to the non-stimulated corresponding control.

### FlaBs mRNA are upregulated in stationary phase

Proteomic and high throughput mass spectrometry performed with the Fiocruz L1-130 strain grown in EMJH have shown that all four *flaBs* genes were expressed and part of the leptospiral flagellum [7]. To test whether leptospires could differently regulate the FlaB expression, cultures of leptospires were harvested after 3 to 6 days, or after 10 to 14 days of culture corresponding to exponential growth or stationary phase, respectively. mRNA was purified and RT-qPCR performed with specific primers of the four leptospiral *flaB* genes. The results suggested that the mRNA expression of the different *flaB* subunits might vary during bacterial growth *in vitro* (Figure 8A). Indeed, the gene expression of all *flaB* subunits seems to be upregulated at the stationary phase in the three serotypes, although only the *flaB3* mRNA expression was statistically higher in stationary phase (Figure 8A). Of note, and different from other strains, the L495 Manilae *flaB1* mRNA was undetectable at the exponential phase, and barely expressed at the stationary phase (Figure 8A). We then compared the human TLR5 activation in presence of the Manilae *flaB1* mutant (the only available *flaB* mutant among pathogenic strains) at the exponential *versus* the stationary phases. In both growth phases, the heat-killed *flaB1* mutant signaled less than the parental Manilae L495 (Figure 8B), confirming that the FlaB1 subunit likely participates in the signaling as already observed (Figure 7C). However, even though the *flaB1* mutant seems to better signal at the stationary phase compared to the exponential phase, no statistical differences could be observed between both phases (Figure 8B). Since in prokaryotes the process from transcription to translation is very rapid, these results of *flaB* mRNA expression together with TLR5 sensing suggest an unanticipated upregulation of the FlaB subunits at the stationary phase or conversely a downregulation at the exponential phase that could potentially influence the TLR5 sensing.

**Figure 8.**
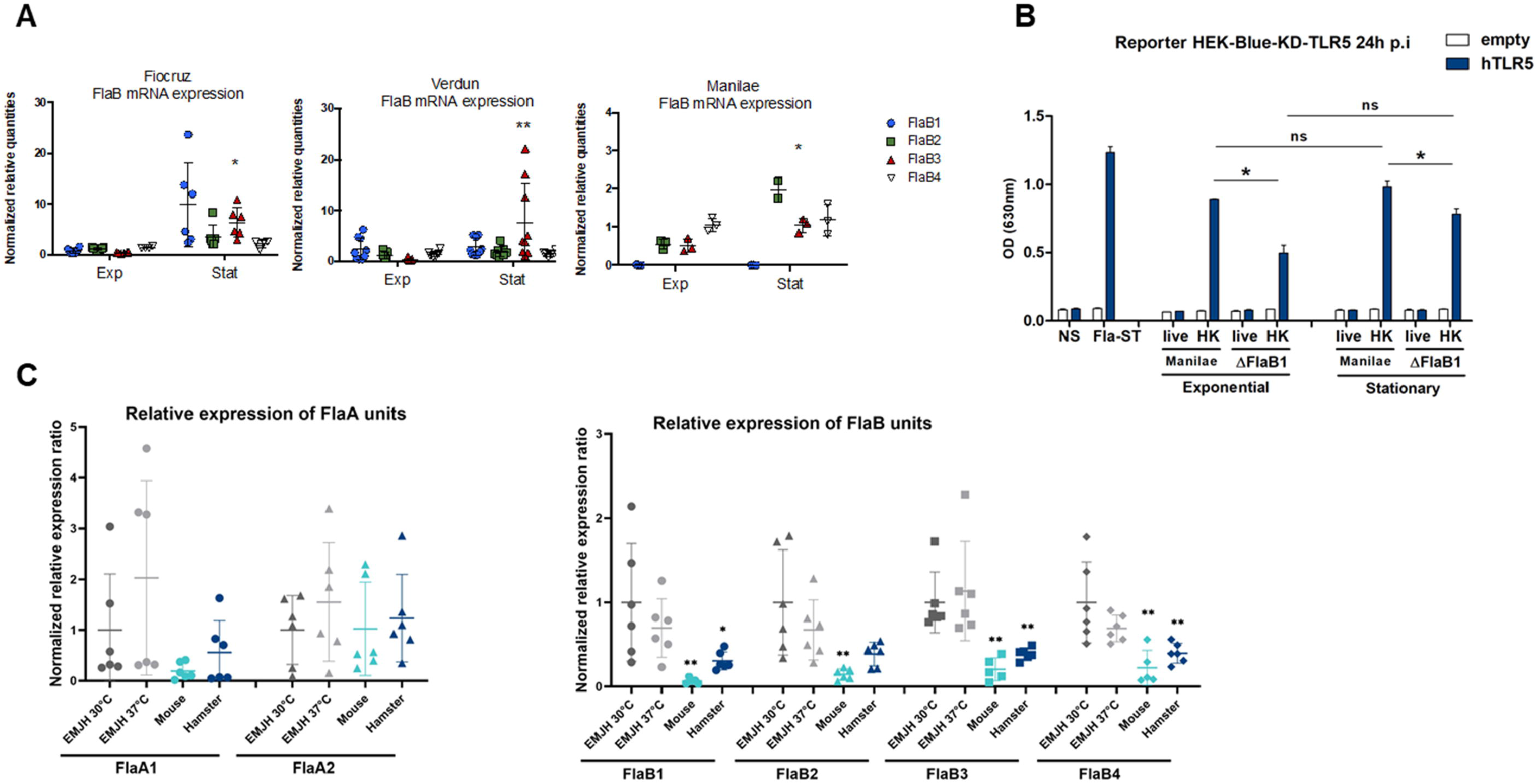
FlaBs mRNA are upregulated in stationary phase and downregulated *in vivo*. **A)** *In vitro* mRNA expression of the different *flaB* subunits in *L. interrogans* (Manilae strain L495, Copenhageni strain Fiocruz L1-130, Icterohaemorrhagiae strain Verdun) at the exponential (exp) and stationary (stat) phase. The RTq-PCR results are expressed as the relative mRNA quantities normalized to the expression of the *lipl41* mRNA. **B)** Reporter NF-κB assay in HEK-Blue-KD-TLR5 cells transfected with the human TLR5 (dark blue bars), or empty plasmid (empty bars) and stimulated with MOI 100 of live or heat-killed *L. interrogans* strain Manilae L495 or ΔFlaB1 mutant in exponential and stationary phase. Flagellin from *Salmonella typhimurium* (Fla-ST) at 500 ng/mL was used as control. Data are expressed as the mean OD630 of n=3 replicates and represent one out of 3 independent experiments and statistics analyses were performed using non-parametric Mann-Whitney test: p<0,05*. **C)** *In vivo* mRNA expression of the different *flaA* and *flaB* subunits in blood of infected mice (n=5) and hamsters (n=5), 24 h post-intraperitoneal infection with 2×10^8^ virulent *L. interrogans* Icterohaemorrhagiae strain Verdun, compared with mRNA expression in EMJH at 30°C and 37°C. Data of RTq-PCR are expressed as the ratio of mRNA quantities relatives to the condition at 30°C. Statistical analyses were performed using non-parametric Mann-Whitney test: p<0,05*. p<0,01**.

### In vivo downregulation of *flaBs* mRNA

To further investigate whether the FlaB regulation could be relevant or play a role *in vivo*, mice and hamsters were infected with the virulent Verdun strain, and blood was sampled 24 hours post infection to purify total mRNAs. In parallel, *in vitro* cultures were performed in EMJH either at 37°C, the host temperature or at 30°C, the usual leptospiral growth culture conditions. First, the expressions of *flaA1* and *flaA2* were not different at 30°C and 37°C, nor between the hamsters and mice, nor between the *in vitro* conditions and *in vivo* conditions (Figure 8C, left panel). However, the *flaB*s expressions were strikingly different, with a weaker expression of the FlaB subunits in the hosts compared to the *in vitro* cultures at 30°C. In the hamsters, all subunits appeared to be equally downregulated compared to their expression in EMJH cultures at 30°C, although in the mouse, FlaB1 seems to be more downregulated than the other subunits, but this was not statistically significant (Figure 8B, right panel). These data strongly suggest that 24h post-infection, leptospires downregulate the expression of their FlaBs subunits in animal blood, which as a consequence could participate in the TLR5 evasion.

## Discussion

We showed in this study that live leptospires largely escaped the TLR5 recognition. However, TLR5 agonists were unexpectedly released after boiling for 30 min, and we further showed their unusual thermoresistance. We determined that the TLR5 activity relied as expected on the FlaB subunits, known to form the core of the flagella and share with FliC some structural features and consensus domains of TLR5 binding. Our results also highlighted a species-specificity of the TLR5 recognition of the leptospiral FlaBs, and potentially differences among strains. Indeed, we evidenced that human and bovine TLR5 recognized heat-killed leptospires, although the mouse TLR5 did not sense the Icterohaemorrhagiae Verdun and Manilae L495 strains, but recognized the Copenhageni Fiocruz strain L1-130, although scantily. We found *in vitro* that the TLR5 recognition was enhanced at the stationary phase, potentially due to an upregulation of the expression of FlaB monomers, especially the FlaB3. Finally, we showed that antimicrobial peptides that are active against live bacteria allowed for their signaling through TLR5. Finally, we showed that leptospires downregulated the FlaBs gene expression in blood from both resistant mice and susceptible hamsters, suggesting a mechanism of immune evasion.

Our results showing the lack of TLR5 signaling by live bacteria despite the involvement of the FlaB subunits in the TLR5 recognition could have been anticipated considering the peculiarities of the leptospiral endoflagella. Indeed, the recent published structure of the filament of the leptospiral flagella showed that the FlaBs form the core and are wrapped inside a lattice composed of both FlaAs, FcpA and FcpB subunits [8], therefore hiding the FlaB monomers. Then, the localization of the flagella inside the bacteria adds a supplemental layer of protection from the host innate immune system. In addition, in *Enterobacteriacea*, a unique FliC monomer polymerizes to form 11 protofilaments that together assemble to constitute one flagellum’s filament. The consensus sites for TLR5 recognition in the flagellin FliC are localized at stacking sites between the flagellin monomers and therefore are not accessible when the filament is formed. Hence, when polymerized, the interaction domain of FliC with TLR5 is masked, therefore whole flagella do not signal through TLR5, [37; 41; 42]. We found that this was also the case for leptospires, with a TLR5 signaling occurring only after 30 min of heating at 70°C, temperature also observed to depolymerize the enterobacterial filament [10]. Similarly, intact purified periplasmic flagella from *Treponema denticola* were not able to activate TLR5 [43].

The absence of TLR5 response in the mouse model was surprising because i) it was shown that neutralizing TLR5 antibodies decreased the cytokine response of whole human blood upon infection with *L. interrogans* [44], ii) we showed here that antimicrobial peptides could degrade live leptospires and induce human TLR5 recognition, and iii) we previously demonstrated that leptospires were killed and cleared from blood during the first days following infection in mice [29], suggesting the release of free flagellin subunits that could have stimulated the TLR5 response. Hence, our study highlights a species-specificity of the TLR5 recognition since murine TLR5, unlike human TLR5, was unable to detect the Manilae L495 and Icterohaemorraghiae Verdun strains. This was unexpected since murine TLR5 is usually more flexible and able to accommodate more different agonist structures compared to human TLR5 [45], similar as seen for murine TLR4 [31]. However, the heat-killed Copenhageni Fiocruz L1-130 strain was recognized by murine TLR5, although to a lesser extent than compared to human TLR5. The weak response seen in murine TLR5 activation is consistent with our previous study showing equivalent levels of IL1ß release in BMMs from WT and TLR5KO mice infected with live Fiocruz L1-130 strain, although stimulation with heat-killed leptospires triggered less IL1ß in TLR5KO BMMs [15]. Interestingly, we previously showed by microdissection of the mouse kidney that TLR5 is expressed in renal tubules, mostly in the distal tubules and in the collecting duct cells while almost not expressed in the proximal tubules [13]. Therefore, our data suggest that the localization of leptospires in proximal tubules could be a favorable environment for the Fiocruz L1-130 strain because of the lack of TLR5 recognition, which potentially could favor its chronic colonization of the mouse kidney [18; 46]. However, since the Manilae L495 strain is not recognized by murine TLR5, we may speculate that it could be advantageous in other animals. Hence, we highlighted an important feature of bovine immune response toward leptospires. Heat killing *Leptospira* revealed their ability to induce a bovine TLR5-dependent response. Antimicrobial peptides also affected live leptospires in a way allowing for the release of TLR5 agonists and subsequent signaling. Of note, bovine antimicrobial peptides are known to have strong potency against leptospires [32]. Hence, we infer that the bovine TLR5 response may be important to fight leptospires in cattle. Together, we may speculate that these observed differences in TLR5 sensing between animals and also between the three strains of *L. interrogans* tested, could, at least partly, be responsible for shaping the preferential species-specificity adaptation of *Leptospira* serovars to their hosts [47].

Interestingly, we also showed a very high stability of the leptospiral filaments and FlaB proteins that perfectly resist heating up to 100°C for 30 min and 85°C for 3 hours. This unusual thermoresistance of the leptospiral flagella is reminiscent the hydrophobic and very highly glycosylated pili of hyperthermophilic Archaea [48]. Glycosylation also occur in bacteria. Although we do not know whether the *Treponema* FlaBs are particularly stable, it has recently been shown that the FlaBs of *Treponema denticola* were glycosylated with an unusual novel glycan [49]. Mass spectrometry analysis of these glycopeptides revealed FlaBs glycosylation by O-linkage at multiple sites near the D1 domain, in the very conserved region of bacterial flagellins that interacts with TLR5 (encompassing the end of consensus 2, Sup Figure 4A)[49]. Interestingly, we found that these atypical glycosylation targets sequences in *Treponema*, notably the two motifs “VEV**S**QL” and “DRIA**S**” are almost 100% conserved in the FlaB1, FlaB2 and FlaB3 of pathogenic and saprophytic *Leptospira* (Sup figure 4B and 4C) [49]. In addition, this consensus was also 100% conserved in leptospiral FlaB1 from other major species involved in leptospirosis in animals and humans (Sup Figure 4D). Interestingly, the two serine residues were substituted in the *L. interrogans* FlaB4 and FlaB from *Borrelia burgdorferi* (Sup Figure 4C), which might suggest a lack of glycosylation of the leptospiral FlaB4 subunit and B. *burdorferi* FLaB. The authors hypothesized that in *Treponema spp*. these peculiar glycosylation could impair the TLR5 signaling of *Treponema*. Our study suggests, if these post-translational modifications exist in leptospires, that they would not impair the TLR5 recognition at least in human and bovine TLR5. Rather we may speculate that they could participate in the thermoresistance of the filament structure.

In the other spirochetes, the filament structure differs from the leptospiral one since in *Treponema* and *Borrelia spp.* the FcpA subunits are absent. Furthermore in *Borrelia* only one copy of FlaA and FlaB compose the filament [50; 51]. The stability of the leptospiral filament is most probably due to the particular association and spatial arrangement of the different FlaBs and to their recently described asymmetric interactions with FlaA and with FcpA [8]. Whether the four FlaBs are randomly dispersed along the filament or would have specific structural functions remains to be studied. However, our results were obtained in the context of the whole bacteria. It would have been interesting to test individual leptospiral FlaB subunits to understand whether the high stability results of intrinsic properties of the individual FlaBs. However, our attempts to express recombinant FlaB monomers have failed. We cannot exclude a caveat in our cloningstrategy but it was quite surprising considering that *T. denticola* and *T. pallidum* FlaB were expressed as stable recombinant proteins that were able to signal through TLR5 in THP1 monocytes or in human keratocytes, respectively [38]. One hypothesis could be that the FlaBs that encompass a different shape than FliC would need to be stabilized by polymerization into the complex filament structure.

The respective role of the leptospiral FlaB1, FlaB2, FlaB3 and FlaB4 proteins remains unknown. The Phyre 2 models suggest that the four FlaBs structures are identical, which explains why the precise roles of the different FlaBs in the core could not be addressed in a recent structural study [8]. The only information available about differences in the four subunits comes from a proteomic study [7] that finds all four FlaB subunits in Fiocruz L1-130 strain cultured in EMJH at 30°C, suggesting that all subunits were present in the filaments with different relative abundance of FlaB subunits, with each bacterium containing 12000 copies of FlaB1, 2000 copies of FlaB2, 300 copies of FlaB3 and 3500 copies of FlaB4 [7]. We tested the expression of each of the four FlaBs mRNA in EMJH cultures and found that all the subunits were expressed in the Verdun and Fiocruz L1-130 strains. However, the relative mRNA levels of the different FlaB subunits did not match the data obtained in the proteomic study, since for example the relative mRNA quantities of *flaB3* seems to be higher than *flaA4* at the stationary phase. Furthermore, in Manilae L495, we observed a strikingly weak expression of FlaB1 compared to the other *L. interrogans* tested, potentially suggesting a strain-specific regulation of FlaB subunits. Of note, the *flaB1* expression was upregulated at the stationary phase in Fiocruz L1-130, which could potentially explain the striking difference between the Manilae L495 strain that was not recognized by the mouse TLR5 whereas the Copenhageni L1-130 strain exhibited a better recognition, despite the fact that all their FlaBs are almost identical and 100% conserved in the TLR5 consensus binding domains. In addition, the absence of one FlaB subunit in the FlaB4 mutant of *L. biflexa* Patoc I, which has been shown to impair the filament formation [19], also impairs the TLR5 signaling. A decreased TLR5 signaling was also observed with the Manilae FlaB1 mutant although the impact of this mutant on filament formation has not yet been studied. However, in both cases the TLR5 signal was not abolished, suggesting that despite the lack of observed motility and filaments, some other FlaB subunits were still expressed and able to signal through TLR5. This is indeed in in agreement with our *in silico* analyzes, suggesting that all FlaB subunits can signal theoretically through TLR5.

The fact that we found a striking downregulation of FlaBs, but not of FlaAs, in the blood of mice and hamsters 24 h post-infection with the Verdun strain, suggests that a regulation of the FlaBs expression could favor an escape from the TLR5 immune surveillance upon infection. However, it remains to be demonstrated that the global downregulation of FlaBs expression that we observed *in vitro* at the exponential phase correlates indeed with a decrease in TLR5 recognition. In animal’s blood, the downregulation of the FlaB expression could make sense to avoid the TLR5 response. It would have been interesting to check the expression of the FlaBs in *Leptospira* colonizing the kidney of animals. However, if amenable in the blood of animals, the purification of leptospires mRNA in kidneys is still challenging. The only example of published renal transcriptome dualseq analysis of *L. interrogans* (Fiocruz L1-130) infection in mice could only detect 29 leptospiral genes [52], among them LipL32, the major lipoprotein and interestingly, one flagellin gene*, flaB4* (LIC11531), suggesting that the mRNA levels of FlaB4 were quite high, and potentially higher than the other FlaBs mRNA. As a whole, these results suggest a complex regulation of the leptospiral FlaB subunits that deserves further investigation. Interestingly, it was shown in another spirochete *Brachyspira hyodysenteriae* that the flagellin genes are transcribed by different transcription factors, with sigma 28 regulating the *flaB1* and *flaB2* genes, whereas the *flaA* and *flaB3* genes are controlled by sigma70. The authors suggest that the relative ratio of the flagellin proteins could play a role in the stiffness of their flagellar filament and consequently that this regulation may play a role in motility [53]. The regulation of FlaBs in leptospires that harbor an even more complex flagellar filament is an interesting question that remains to be studied.

Interestingly, the leptospiral FlaBs share with the flagellin of *Bacillus spp*, that is also able to signal via TLR5, a similar structure made of the D0 and D1 domains of FliC and lacking the D2 and D3 domains [54]. Of note, the D2 and D3 domains of FliC are highly variable and responsible for the strong antigenicity of flagellins in *Enterobacteriacae* [10]. Flagellin is known to be a potent vaccine adjuvant, however the antigenicity of the D2 and D3 domains can be a problem when booster immunizations are done. To circumvent this issue, several strategies have been recently proposed. The first consisted in using a FliC devoid of the D2 and D3 domains [14], and the second to use the *Bacillus* flagellin as an expression platform [54]. Likewise, we may speculate in the case of *Leptospira spp* that upon *in vivo* killing and exposure of FlaB subunits, the lack of D2 and D3 domains could be advantageous to limit the antibody response. Hence, the peculiar structure of FlaBs could also participate in the adaptive immune evasion

In conclusion, we showed here that pathogenic *Leptospira* largely escape recognition by TLR5. Other bacteria such as *Helicobacter pylori* have been shown to escape the TLR5 response through modification of the amino residues in the D0 or D1 regions of flagellin subunits [39], but leptospires seem to have developed a different escape strategy. Indeed, our data demonstrate that the endoflagella play a role in the escape from TLR5 surveillance, which has never been shown before and might hold true for other spirochetes. We also evidenced regulatory mechanisms of *FlaB* genes expression that may also play a role in this immune evasion and have important consequences since TLR5 ligation has a potent adjuvant role in immunity.

## Acknowledgements

We thank Brigitte David-Watine for critical reading of the manuscript. We are grateful to Marie-Estelle Soupe-Gilbert for her participation in the design of primers used for the quantification of flagellar subunits gene expression.

This study received funding from the French Government’s Investissement d’Avenir program, Laboratoire d’Excellence “Integrative Biology of Emerging Infectious Diseases” (grant n°ANR-10-LABX-62-IBEID) to IGB. DB received funding from the Ecole Doctorale Frontières de l’Innovation en Recherche et Education (FIRE), Programme Bettencourt. JC was supported by a Calmette and Yersin fellowship from Institut Pasteur International network and EW by a NIH fund (R01AI121207).

