## Supplemental figures for "Escape of TLR5 Recognition by *Leptospira spp*: A Rationale for Atypical Endoflagella"

### Slide 1
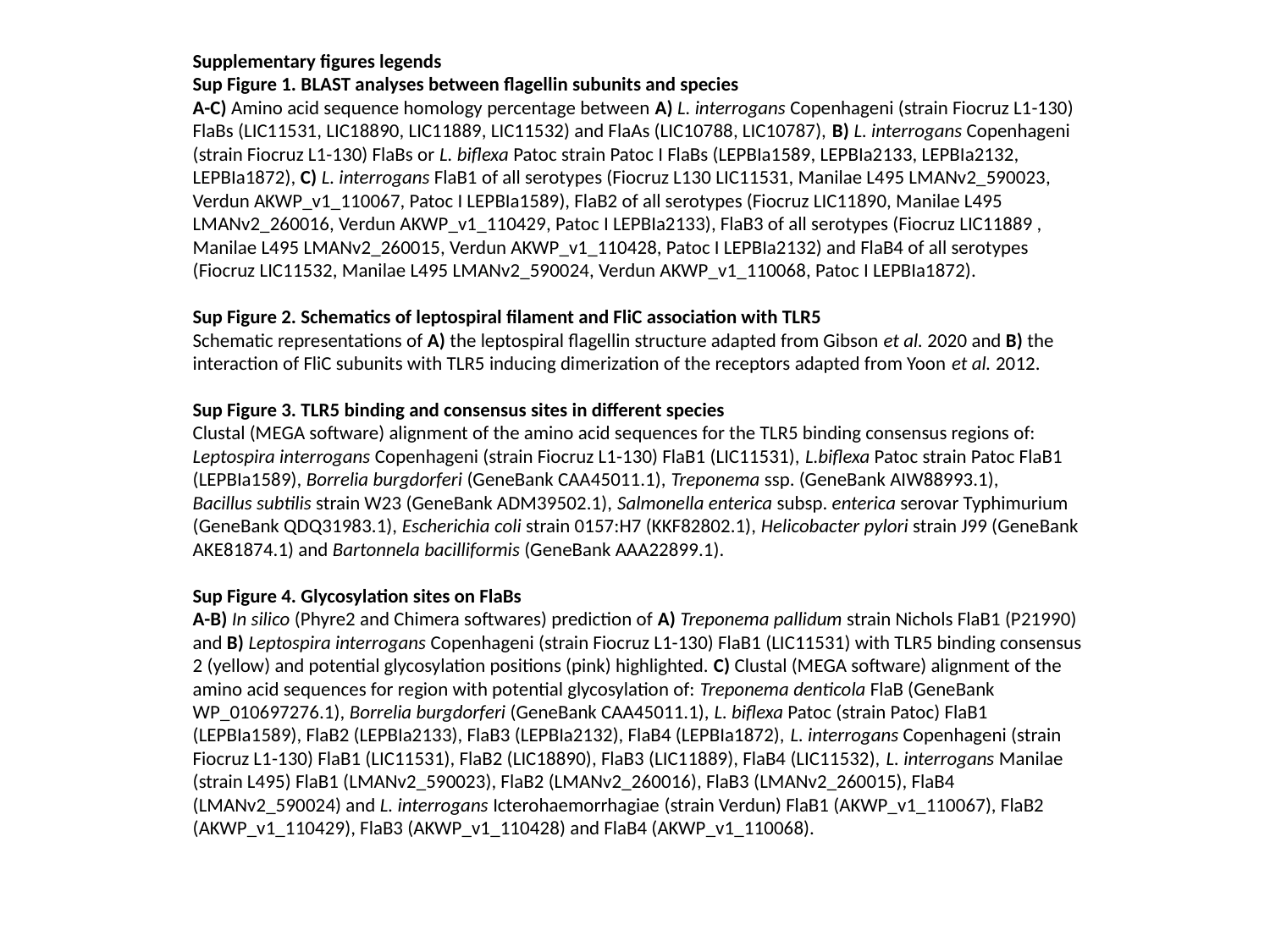

Supplementary figures legends
Sup Figure 1. BLAST analyses between flagellin subunits and species
A-C) Amino acid sequence homology percentage between A) L. interrogans Copenhageni (strain Fiocruz L1-130) FlaBs (LIC11531, LIC18890, LIC11889, LIC11532) and FlaAs (LIC10788, LIC10787), B) L. interrogans Copenhageni (strain Fiocruz L1-130) FlaBs or L. biflexa Patoc strain Patoc I FlaBs (LEPBIa1589, LEPBIa2133, LEPBIa2132, LEPBIa1872), C) L. interrogans FlaB1 of all serotypes (Fiocruz L130 LIC11531, Manilae L495 LMANv2_590023, Verdun AKWP_v1_110067, Patoc I LEPBIa1589), FlaB2 of all serotypes (Fiocruz LIC11890, Manilae L495 LMANv2_260016, Verdun AKWP_v1_110429, Patoc I LEPBIa2133), FlaB3 of all serotypes (Fiocruz LIC11889 , Manilae L495 LMANv2_260015, Verdun AKWP_v1_110428, Patoc I LEPBIa2132) and FlaB4 of all serotypes (Fiocruz LIC11532, Manilae L495 LMANv2_590024, Verdun AKWP_v1_110068, Patoc I LEPBIa1872).
Sup Figure 2. Schematics of leptospiral filament and FliC association with TLR5
Schematic representations of A) the leptospiral flagellin structure adapted from Gibson et al. 2020 and B) the interaction of FliC subunits with TLR5 inducing dimerization of the receptors adapted from Yoon et al. 2012.
Sup Figure 3. TLR5 binding and consensus sites in different species
Clustal (MEGA software) alignment of the amino acid sequences for the TLR5 binding consensus regions of: Leptospira interrogans Copenhageni (strain Fiocruz L1-130) FlaB1 (LIC11531), L.biflexa Patoc strain Patoc FlaB1 (LEPBIa1589), Borrelia burgdorferi (GeneBank CAA45011.1), Treponema ssp. (GeneBank AIW88993.1), Bacillus subtilis strain W23 (GeneBank ADM39502.1), Salmonella enterica subsp. enterica serovar Typhimurium (GeneBank QDQ31983.1), Escherichia coli strain 0157:H7 (KKF82802.1), Helicobacter pylori strain J99 (GeneBank AKE81874.1) and Bartonnela bacilliformis (GeneBank AAA22899.1).
Sup Figure 4. Glycosylation sites on FlaBs
A-B) In silico (Phyre2 and Chimera softwares) prediction of A) Treponema pallidum strain Nichols FlaB1 (P21990) and B) Leptospira interrogans Copenhageni (strain Fiocruz L1-130) FlaB1 (LIC11531) with TLR5 binding consensus 2 (yellow) and potential glycosylation positions (pink) highlighted. C) Clustal (MEGA software) alignment of the amino acid sequences for region with potential glycosylation of: Treponema denticola FlaB (GeneBank WP_010697276.1), Borrelia burgdorferi (GeneBank CAA45011.1), L. biflexa Patoc (strain Patoc) FlaB1 (LEPBIa1589), FlaB2 (LEPBIa2133), FlaB3 (LEPBIa2132), FlaB4 (LEPBIa1872), L. interrogans Copenhageni (strain Fiocruz L1-130) FlaB1 (LIC11531), FlaB2 (LIC18890), FlaB3 (LIC11889), FlaB4 (LIC11532), L. interrogans Manilae (strain L495) FlaB1 (LMANv2_590023), FlaB2 (LMANv2_260016), FlaB3 (LMANv2_260015), FlaB4 (LMANv2_590024) and L. interrogans Icterohaemorrhagiae (strain Verdun) FlaB1 (AKWP_v1_110067), FlaB2 (AKWP_v1_110429), FlaB3 (AKWP_v1_110428) and FlaB4 (AKWP_v1_110068).

### Slide 2
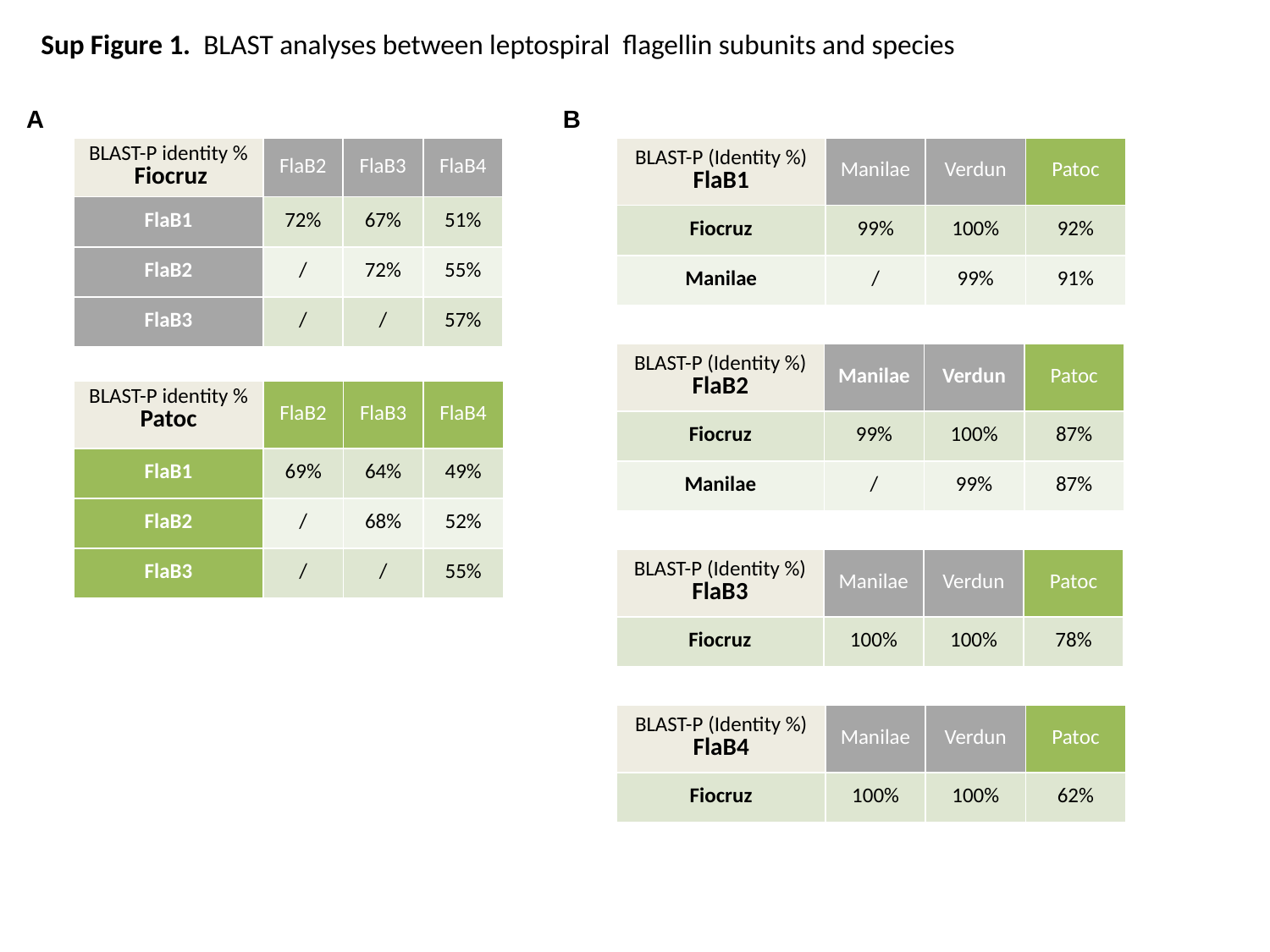

Sup Figure 1. BLAST analyses between leptospiral flagellin subunits and species
A
B
| BLAST-P (Identity %) FlaB1 | Manilae | Verdun | Patoc |
| --- | --- | --- | --- |
| Fiocruz | 99% | 100% | 92% |
| Manilae | / | 99% | 91% |
| BLAST-P identity % Fiocruz | FlaB2 | FlaB3 | FlaB4 |
| --- | --- | --- | --- |
| FlaB1 | 72% | 67% | 51% |
| FlaB2 | / | 72% | 55% |
| FlaB3 | / | / | 57% |
| BLAST-P (Identity %) FlaB2 | Manilae | Verdun | Patoc |
| --- | --- | --- | --- |
| Fiocruz | 99% | 100% | 87% |
| Manilae | / | 99% | 87% |
| BLAST-P identity % Patoc | FlaB2 | FlaB3 | FlaB4 |
| --- | --- | --- | --- |
| FlaB1 | 69% | 64% | 49% |
| FlaB2 | / | 68% | 52% |
| FlaB3 | / | / | 55% |
| BLAST-P (Identity %) FlaB3 | Manilae | Verdun | Patoc |
| --- | --- | --- | --- |
| Fiocruz | 100% | 100% | 78% |
| BLAST-P (Identity %) FlaB4 | Manilae | Verdun | Patoc |
| --- | --- | --- | --- |
| Fiocruz | 100% | 100% | 62% |

### Slide 3
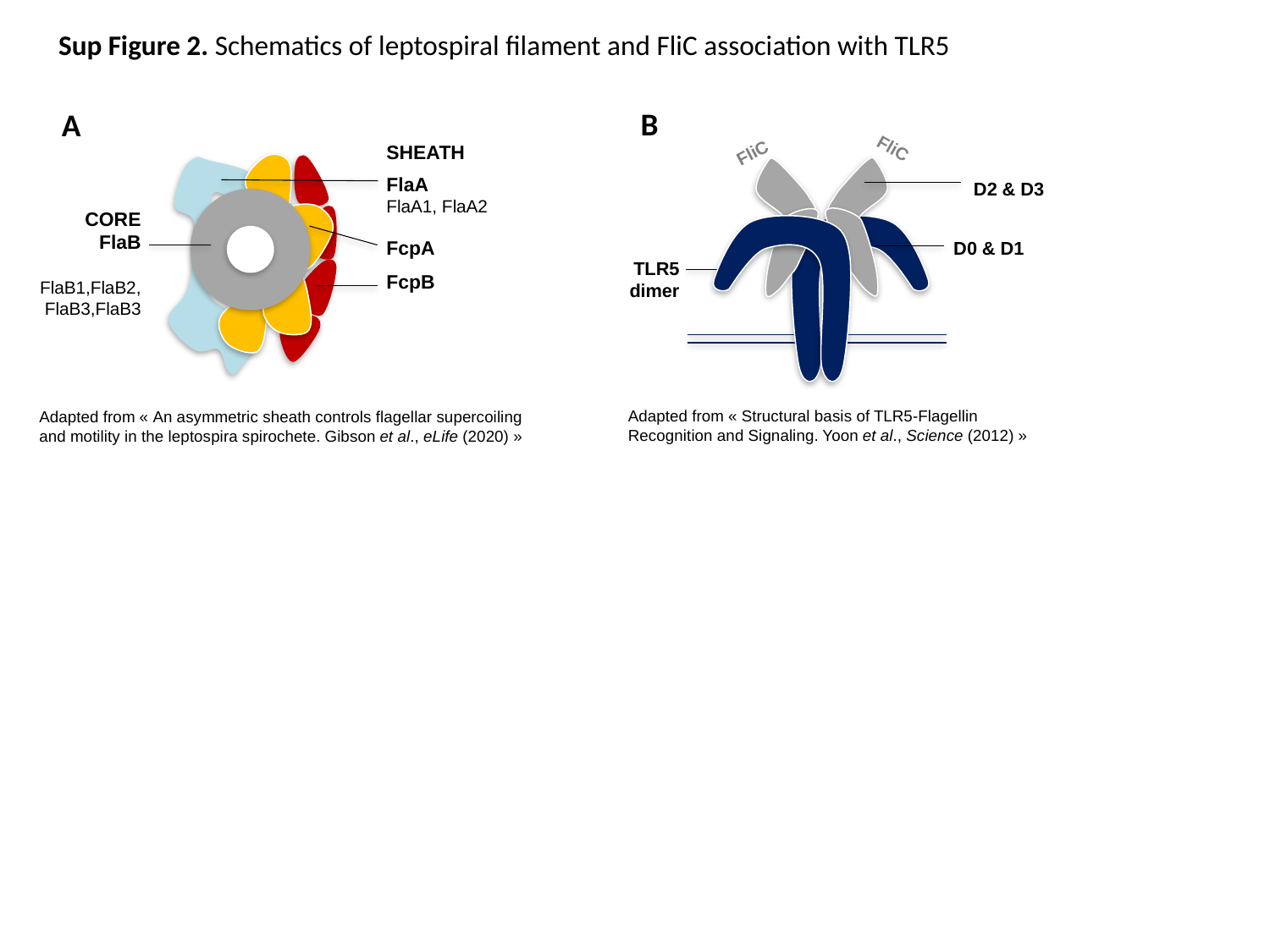

Sup Figure 2. Schematics of leptospiral filament and FliC association with TLR5
B
A
FliC
SHEATH
FlaA
FlaA1, FlaA2
FcpA
FcpB
CORE
FlaB
FlaB1,FlaB2,
FlaB3,FlaB3
FliC
D2 & D3
D0 & D1
TLR5
dimer
Adapted from « Structural basis of TLR5-Flagellin Recognition and Signaling. Yoon et al., Science (2012) »
Adapted from « An asymmetric sheath controls flagellar supercoiling and motility in the leptospira spirochete. Gibson et al., eLife (2020) »

### Slide 4
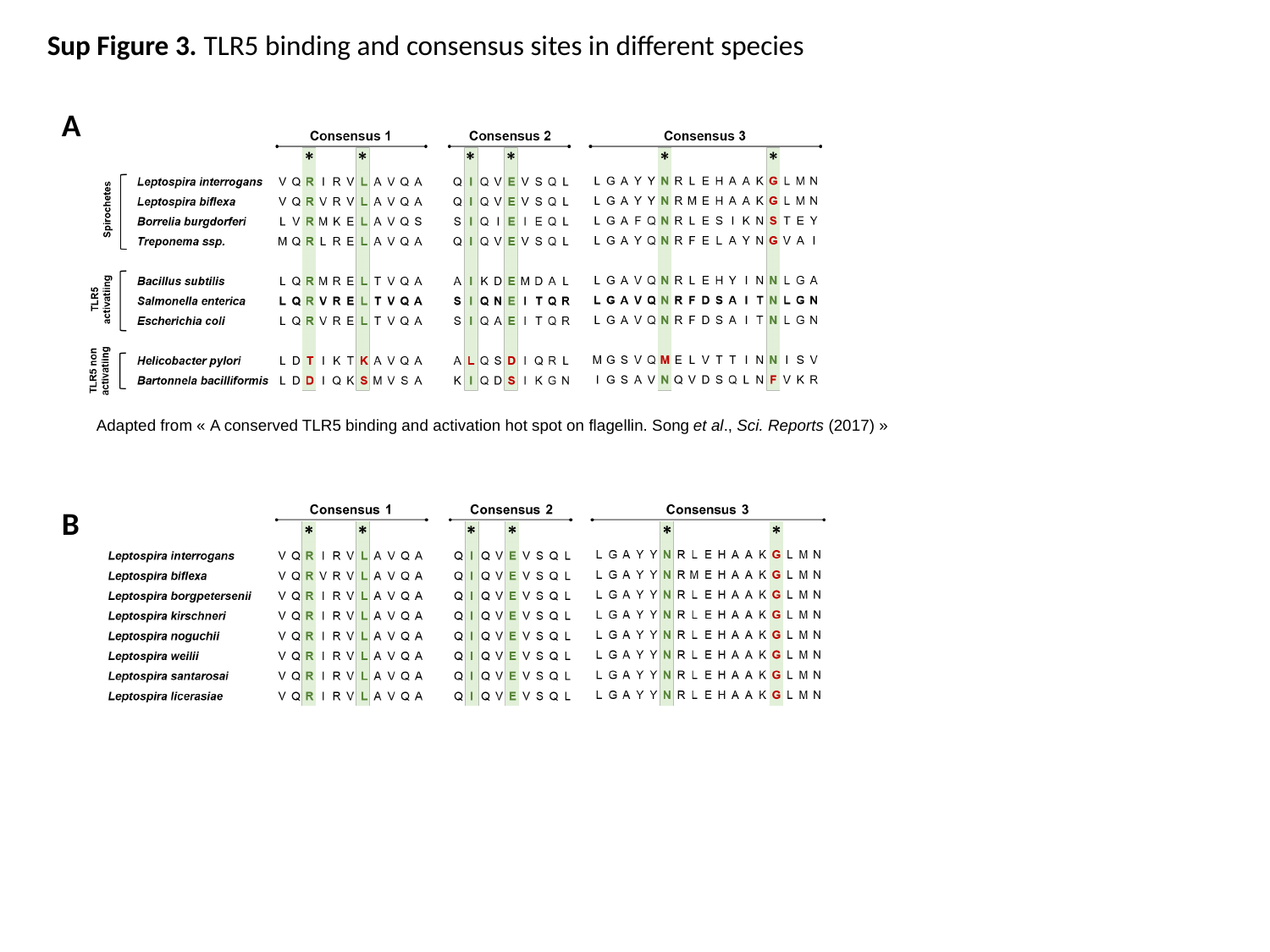

Sup Figure 3. TLR5 binding and consensus sites in different species
A
Adapted from « A conserved TLR5 binding and activation hot spot on flagellin. Song et al., Sci. Reports (2017) »
B

### Slide 5
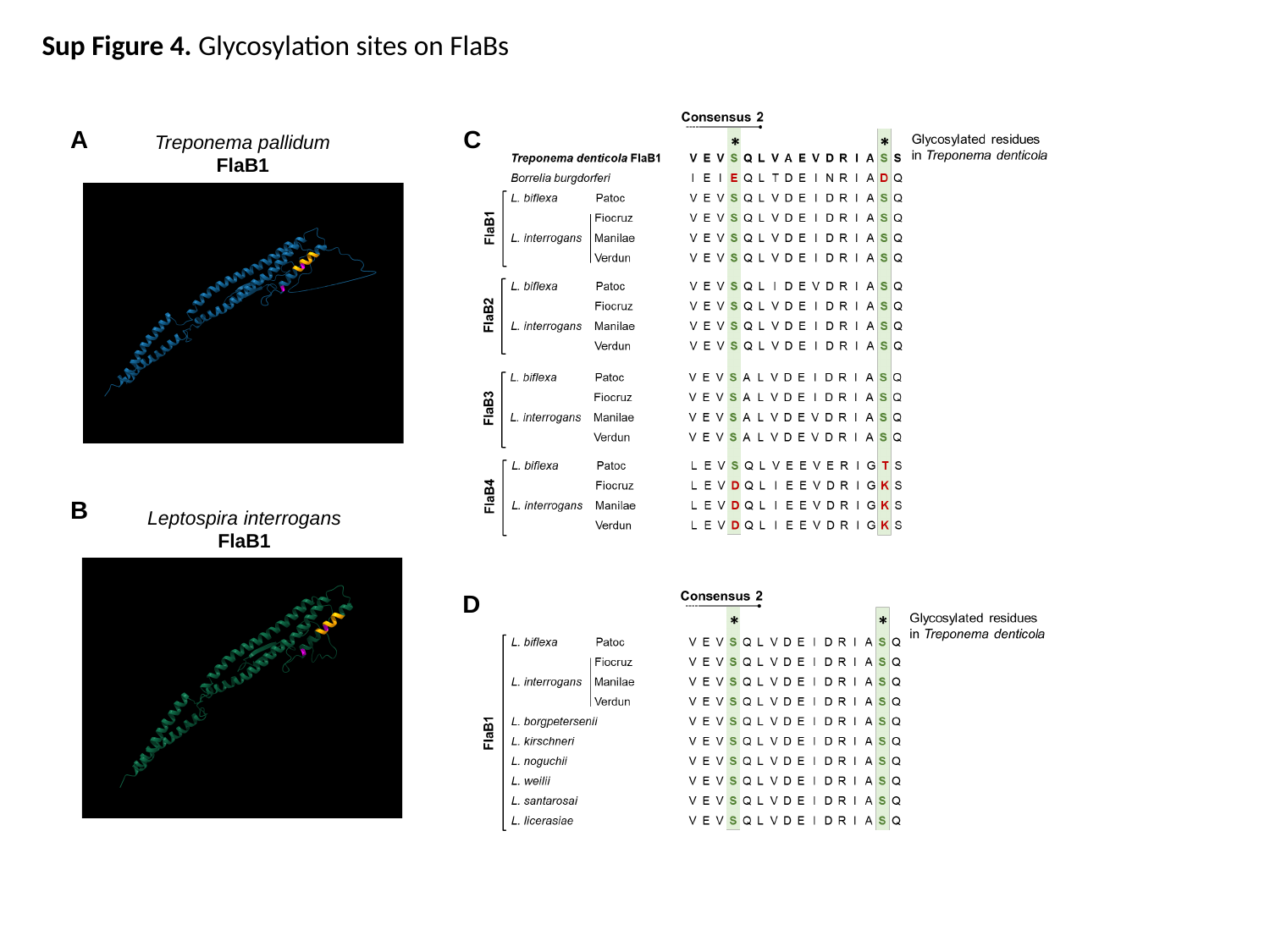

Sup Figure 4. Glycosylation sites on FlaBs
A
C
Treponema pallidum
FlaB1
B
Leptospira interrogans
FlaB1
D
